# A minimally modified AAV9 variant (AAV9fyhr) for enhanced CNS gene delivery by intra-CSF administration

**DOI:** 10.64898/2026.10.07.756773

**Authors:** Irvin T. Garza, Vijayashree Ramesh, Yuhui Hu, Matt Rioux, Mehmet Takar, Harrison Higgs, Erik A. Lykken, Rachel Adams, Peter Kunach, Daniel Kieffer, Cynthia B Trygg, Jeanne M. Fisher-Perkins, Ashwani Kumar, Lukasz A Joachimiak, Rachel M. Bailey, Chao Xing, Stephen E. Braun, Steven J. Gray

## Abstract

Adeno-associated virus (AAV) mediated gene therapies continue to show promise for the treatment of genetic diseases. However, delivery and safety still represent significant prohibitive barriers restricting translation of these delivery modalities to humans. We describe a novel AAV9 variant, “AAV9fyhr” or “FYHR”, identified from screening a controlled randomization point mutation AAV9 library in mice and nonhuman primates followed by directed evolution of in vivo CNS-derived libraries screened in human iPSC differentiated cortical neurons. Capsid variants of interest were identified by coupling long read sequencing and computational structure modeling. Structure-function analysis of our lead candidate reveal that two targeted amino acid changes are sufficient to result in beneficial pleiotropic effects across viral stability, cell surface attachment, and gene delivery. We developed a novel multiplexed barcoding reporter system, enabling a time and cost-effective option to gather spatially resolved single cell capsid validation data in mice and nonhuman primates. When administered intrathecally, AAV9fyhr exhibits 2-to-6-fold higher gene delivery across several tissues in the CNS while minimally altering the tropism to peripheral tissues in comparison to AAV9. These findings demonstrate the potential of whole capsid engineering for the rational development of translational next generation AAV capsids.

## INTRODUCTION

The modern molecular biology toolkit has enabled the potential to develop precision medicine, or gene therapies, that address the underlying cause of thousands of genetic diseases. Gene therapies act by delivering entire genes^1,2^, silencing^3–6^ or attenuating unwanted genetic information; and even correcting or editing host genomic sequences^7–9^ *ex vivo* and *in vivo*. While recent gene therapies have shown promising results^10,11^, scalability^12,13^, efficacy^14,15^, and safety^16–19^ remain as the primary prohibitive barriers for future applications. All these issues can be positively impacted with improved delivery technology.

Adeno-associated viral vectors are small non-pathogenic delivery vehicles that have demonstrated a relatively safe performance history for the long-lasting single administration delivery of DNA cargos to multiple tissues. Although they have a limited packaging capacity, multiple strategies have been implemented for the delivery of larger genes, silencing agents, and gene editing machinery^20–23^. Currently, there are hundreds of ongoing clinical trials making use of naturally occurring AAVs for a variety of tissues, including challenging extrahepatic tissues such as the brain, muscle, and eye^1,24,25^. Recently, the technology in AAV capsids has seen a tremendous improvement in modifications that have improved diversification strategies, targeted delivery, and liver detargeting^26–30^. Much of these strategies take advantage of surface exposed loops that are amenable to peptide insertions, and while these strategies have yielded promising results, these modifications may have possible effects on capsid assembly which ultimately may pose a challenging barrier in vector manufacturing^25,29,31–33^. Furthermore, many of these strategies utilize libraries of peptide insertions into naturally occurring AAV capsids, such as AAV9. Historically, sequencing technology limited the ability to generate and evaluate complex libraries at sufficient throughput. However, advances in long-read sequencing technology have made whole capsid engineering increasingly viable^34–36^. This approach enables interrogation of interactions between residues that may be distant in the linear amino acid sequence but become spatially proximal within the three-dimensional capsid structure, potentially revealing cooperative effects that influence capsid function. Whole capsid engineering therefore has the potential to enhance multiple AAV properties, yielding improved capsids for gene therapy and may serve as novel base capsids that can be coupled with other capsid modification strategies, such as peptide insertions.

To undertake this, we generated a novel AAV9 controlled random mutagenesis library. This strategy introduces mutations throughout the capsid sequence at a controlled frequency and sequence space, enabling interrogation of capsid sequence space while retaining sufficient similarity to the parental AAV9 scaffold for functional selection. Upon library generation, capsids were screened in parallel within mice and nonhuman primates by intrathecal lumbar cistern administration to identify species cross-compatible capsids. Capsid variants were recovered from nuclei isolated out of CNS tissues, increasing selection stringency for capsids capable of successfully delivering their genetic payload to the nucleus^35^. Four lead candidates were identified and further characterized using a multiplexed reported capsid validation system alongside AAV9, serving as a benchmark. Our lead capsid candidate, AAV9fyhr and herein referred to as “FYHR”, demonstrates that two amino acid mutations on the AAV9 sequence are sufficient to increase gene delivery both in-vitro and in-vivo. Characterization of FYHR suggest a pleiotropic effect that culminates in overall enhancement mediated through multiple viral transduction checkpoints. These findings provide a fundamental understanding of AAV9 capsid biology that facilitates a whole capsid engineering strategy with accelerated capsid validation for translational gene therapy applications.

## RESULTS

### Identification of AAV9fyhr by screening a controlled randomization AAV9 library in mice and nonhuman primates

AAVs have 2 major open reading frames (ORFs), known as Rep and Cap. Cap encodes all of the structural proteins that assemble into the capsid, however, two accessory proteins, AAP^37^ and MAAP^38^, lie in alternative open reading frames within the Cap gene playing a critical role in viral assembly and viral egress, respectively. More specifically, AAV9 has shown to have broad tissue tropism with blood-brain barrier crossing capacity. For these reasons, we designed a controlled randomization library of AAV9 variants that focused only on variable regions III-IX, leaving the remainder of the capsid unaltered (Fig. 1A). This approach does not alter AAP or MAAP, which do not overlap with variable regions III-IX. The library uses a wildtype AAV plasmid vector providing the AAV inverted terminal repeats (ITRs) as well as Rep and Cap genes in *cis*. AAVs are 60mer biomolecular complexes that assemble into pseudo symmetrical structures. Therefore, high throughput capsid library generation strategies require careful library production strategies to ensure that AAV variants are not predominantly mosaic assemblies of multiple individual Cap subunit variants. To generate the library, we first packaged the genomes of variants into an AAV2 capsid, herein referred to as our “shuttle” library. The purified shuttle vector containing the packaged AAV9 variants is then subsequently co-infected in a second round of production with wildtype adenovirus (Ad5) such that a single AAV particle infects a single cell (Fig. 1A and Methods). To select for species cross-compatible variants with enriched gene delivery to the CNS, adult mice and juvenile nonhuman primates were injected in parallel via the intrathecal lumbar cistern. Two weeks later, AAV9 variants that successfully trafficked to the nuclei were recovered from brain and spinal cord samples by isolating DNA from nuclear fractions. To identify variants, we utilized single molecule real time (SMRT) sequencing on PCR recovered amplicons of the full 2.2 kb cap gene (Fig. 1B). After one round of selection *in vivo*, AAV9fyhr, herein referred to as “FYHR”, showed relatively similar enrichment in mice and NHPs across several samples in the CNS when compared to AAV9 (Fig. 1C).

**Figure 1.**
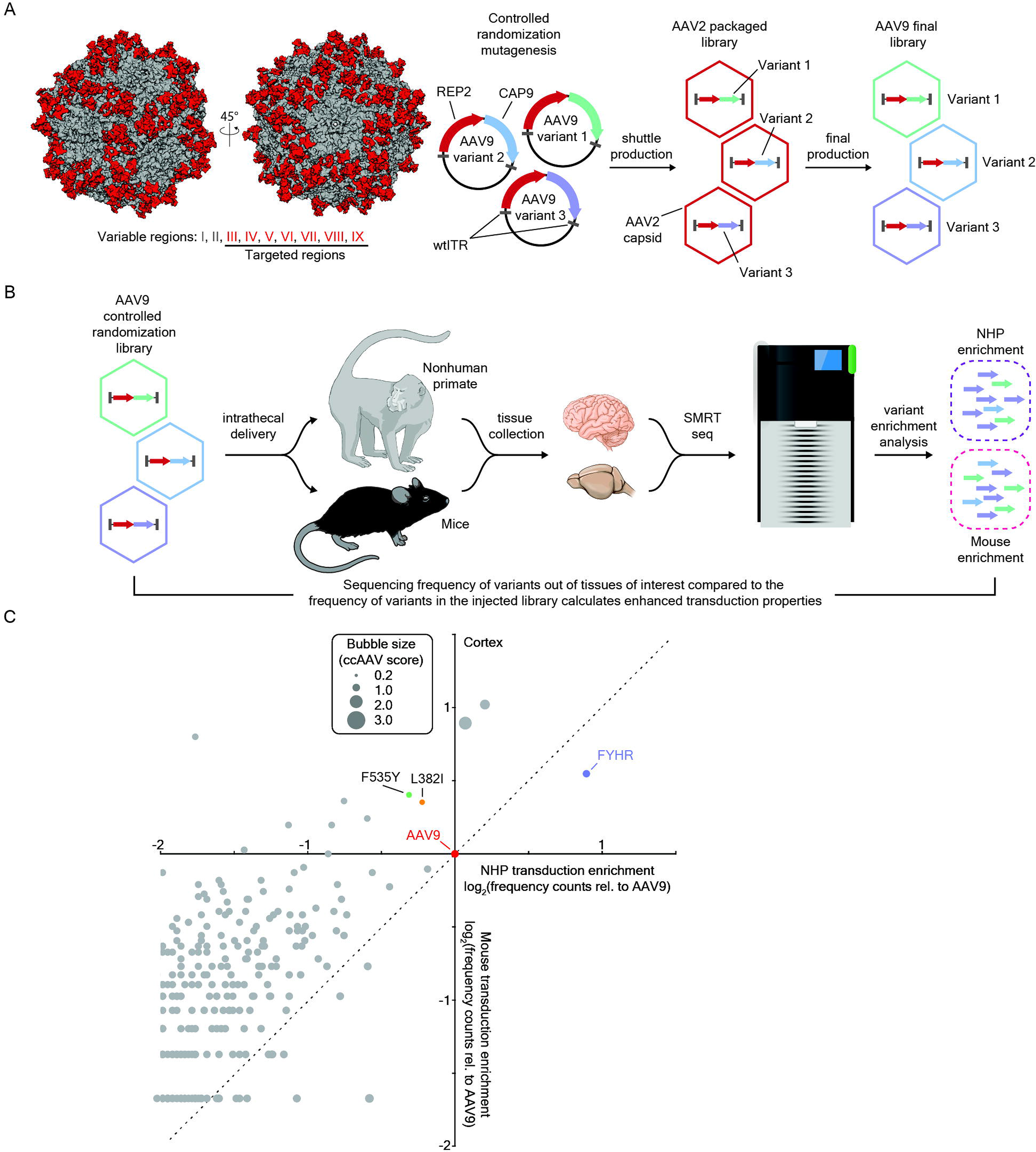
In vivo screens of a controlled randomization library of AAV9. (A) Location of controlled randomization regions (red) on the AAV capsid shown on the structure of AAV9 (PDB: 7MT0)^48^ and schematic of AAV library generation. (B) Schematic of parallel AAV library screening administered by a lumbar intrathecal route in mice and nonhuman primates. Variants are recovered from isolated nuclei from CNS tissues and identified by long read sequencing. (C) Scatterplot of log_2_ frequency counts of recovered variant from cortex of NHP (x-axis) and mice (y-axis) are displayed relative to AAV9. Individual variants are represented as a single bubble. Bubble size corresponds to the ratio of mouse to NHP log_2_ frequency count relative to AAV9 of each variant. Dashed line represents a slope = 1.

### Directed evolution of CNS derived variants in human iPSC differentiated cortical neurons refine selections

AAV9 variants recovered from mice and NHP’s CNS tissues were then used to generate a second round of tissue-derived libraries. To identify capsids amenable to human neuronal gene delivery, both mice and NHP CNS-derived libraries were applied to human induced pluripotent stem cell (hiPSC) derived cortical neurons (Fig. 2A). Capsids were recovered and identified as described in the mouse and NHP screens. Variant enrichment analysis of two replicates demonstrated that FYHR and H584R were the most prominent capsid variants recovered (Fig. 2B). Furthermore, a second variant, AAV9hrli (“HRLI”), and several other variants also incorporated this H584R mutation within our screen.

**Figure 2.**
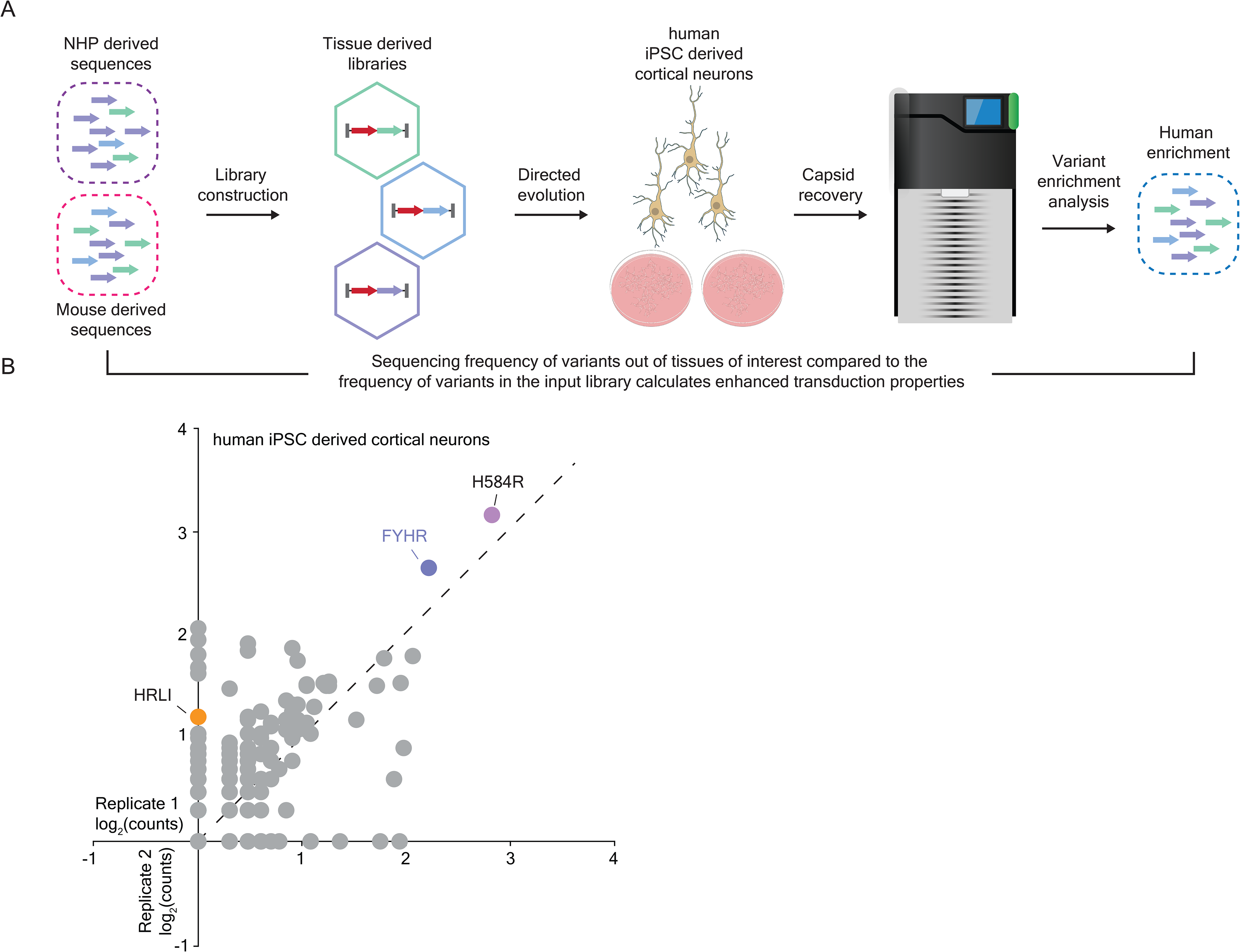
Directed evolution in human iPSC differentiated neurons. (A) Schematic of CNS recovered variants from mice and NHP screened through iPSC derived cortical neurons. (B) Scatterplot of log_2_ frequency counts of recovered variants in biological replicates (n = 2). Dashed line represents a slope = 1.

### Sequence and structural analysis of AAV9fyhr amino acid changes

AAV9fyhr was selected as our lead candidate for further investigation and validation. Compared with wild-type AAV9, AAV9fyhr harbors two amino acid substitutions, F535Y and H584R. To begin defining the potential contributions of each mutation, we performed sequence and structural alignments (Table 1, Fig. 3A & 3B) across common naturally occurring AAV serotypes (AAV1-13). Position 535 was highly conserved, with all serotypes retaining a phenylalanine except the evolutionarily divergent AAV5, which contains the similarly hydrophobic leucine residue (Table 1). Structurally, residue 535 is relatively buried within the viral protein core (Figure S1), suggesting that F535Y may primarily influence local capsid structure or biophysical properties. In contrast, position 584 exhibited substantially greater sequence diversity. AAV9 contains a histidine at this position, whereas most naturally occurring serotypes contain hydrophobic or uncharged residues, with AAV4 containing a negatively charged aspartic acid (Table 1). Given its greater surface exposure (Figure S1), we hypothesized that H584R may be the primary driver of altered interactions between the viral capsid and host-cell factors.

**Figure 3.**
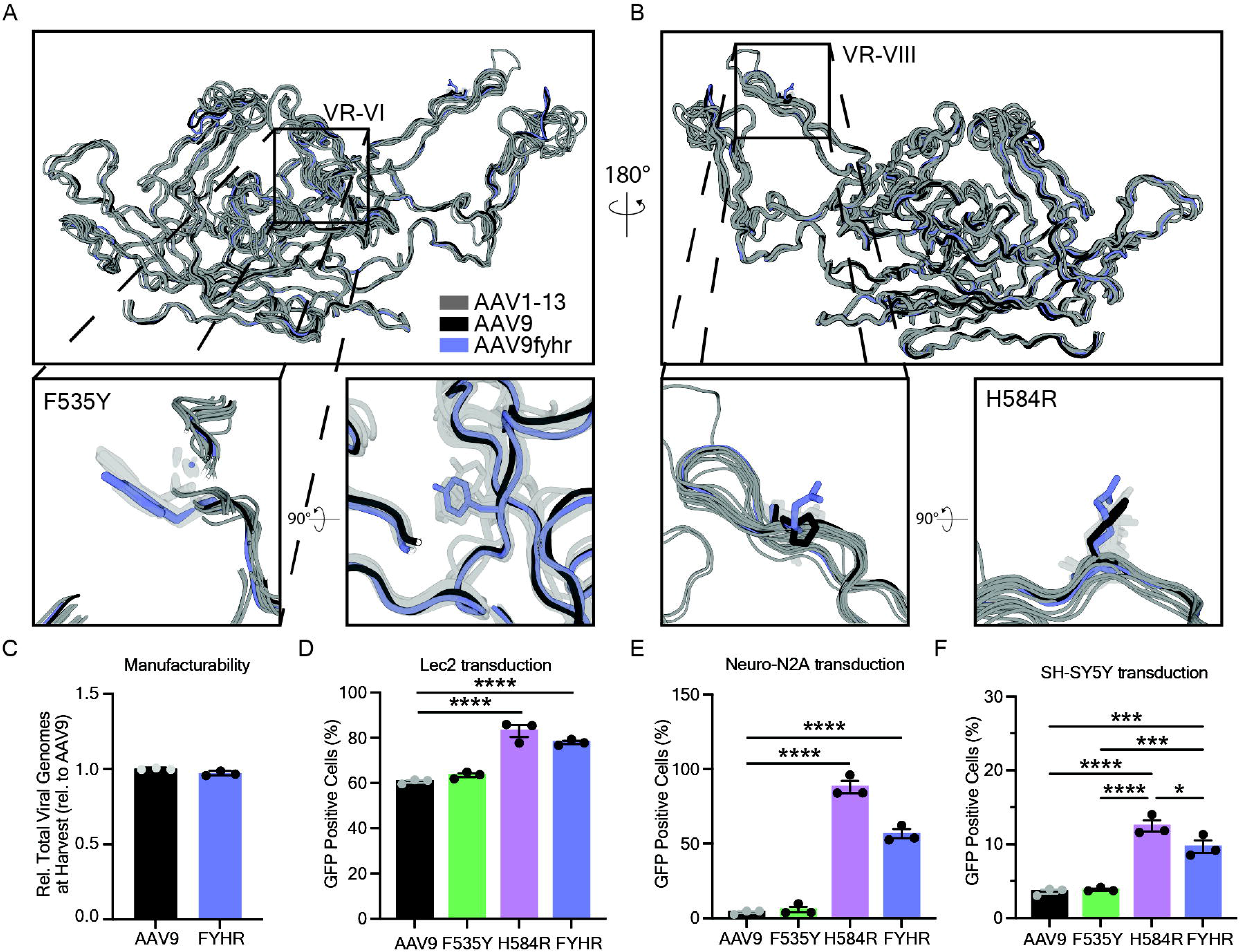
In vitro characterization of AAV9 variants reveals that H584R is sufficient for AAV9 transduction enhancement. (A) Structural alignments of AAV serotypes from Table 1 to a Cryo-EM structure of AAV9fyhr comparing the residue at 535 (rel. to AAV9 VP1) on VP3. (B) A 180° rotational view to compare the residues at 584 (rel. AAV9 VP1). (C) Comparison of recombinant AAV9 or AAV9fyhr production yields. (D-F) Transduction assessed by flow cytometry of CHO Lec2 (D), Neuro-N2A (E), and SH-SY5Y (F). Manufacturability data points represent mean values from three independent experiments of technical triplicates; transduction data represents n = 3 well replicates. Student’s t-test was used for manufacturability; One-way ANOVA was used with Sidak multiple comparison correction for transduction comparisons. Error bars indicate ± SD. *p<0.05, ***p<0.001, ****p<0.0001

**Table 1.** Comparison of F535 and H584 amino acids and chemical property across most naturally occurring AAV serotypes. Capsid, serotype; VP1 aa, VP1 sequence length, 535 aa rel. AAV9, structurally identical residue relative to AAV9; Chemical property at 535, chemical property of corresponding residue at 535 equivalent relative to AAV9; 584 aa rel. AAV9, structurally identical residue relative to AAV9; Chemical property at 584, chemical property of corresponding residue at 584 equivalent relative to AAV9.

| Capsid | VP1 aa | 535 aa<br>rel. AAV9 | Chemical<br>Property at 535 | 584 aa<br>rel. AAV9 | Chemical<br>Property at 584 |
| --- | --- | --- | --- | --- | --- |
| FYHR | 736 | Tyrosine (Y) | Mildly hydrophilic | Arginine (R) | Positively Charged |
| AAV1 | 736 | Phylalanine (F) | Hydrophobic | Phylalanine (F) | Hydrophobic |
| AAV2 | 735 | Phylalanine (F) | Hydrophobic | Leucine (L) | Hydrophobic |
| AAV3B | 735 | Phylalanine (F) | Hydrophobic | Leucine (L) | Hydrophobic |
| AAV4 | 735 | Phylalanine (F) | Hydrophobic | Aspartic Acid (D) | Negatively Charged |
| AAV5 | 736 | Leucine (L) | Hydrophobic | Asparagine (N) | Polar No Charge |
| AAV6 | 736 | Phylalanine (F) | Hydrophobic | Leucine (L) | Hydrophobic |
| AAV7 | 735 | Phylalanine (F) | Hydrophobic | Leucine (L) | Hydrophobic |
| AAV8 | 735 | Phylalanine (F) | Hydrophobic | Leucine (L) | Hydrophobic |
| AAV9 | 736 | Phylalanine (F) | Hydrophobic | Histidine (H) | Positively Charged |
| AAVrh10 | 736 | Phylalanine (F) | Hydrophobic | Leucine (L) | Hydrophobic |
| AAV11 | 735 | Phylalanine (F) | Hydrophobic | Asparagine (N) | Polar No Charge |
| AAV12 | 735 | Phylalanine (F) | Hydrophobic | Asparagine (N) | Polar No Charge |
| AAV13 | 735 | Phylalanine (F) | Hydrophobic | Asparagine (N) | Polar No Charge |

### H584R is sufficient to enhance transduction without compromising vector production

To empirically determine the individual contributions of these mutations, we generated recombinant AAV9, F535Y, H584R, and FYHR vectors carrying a GFP reporter transgene. FYHR displayed vector yields comparable to AAV9, indicating that the combined mutations did not measurably impair manufacturability (Fig. 3C). We next asked whether enhanced transduction resulted from altered utilization of AAV9’s canonical terminal galactose receptor. Chinese hamster ovary (CHO) lec2 cells, which display increased cell-surface galactose, were transduced with each vector. H584R and FYHR exhibited increased transduction relative to AAV9, whereas F535Y produced only a modest, non-significant increase (Fig. 3D). We similarly assessed whether the variants remained responsive to terminal galactose availability. Treatment of HEK293 cells with neuraminidase, which increases exposure of cell-surface galactose by cleavage of terminal sialic acid on the cell surface, enhanced AAV9 transduction as expected and produced comparable increases in transduction for F535Y, H584R, and FYHR (Figure S2.). These results indicate that H584R is sufficient to reproduce much of the enhanced transduction phenotype observed with FYHR while preserving the parent AAV9 productive entry through a galactose-rich cellular context.

### H584R enhances transduction across multiple cell types

We next asked whether the effect of H584R was restricted to CHO lec2 and HEK293T cells or represented a broader enhancement in cellular transduction. In both mouse (Neuro-N2A) and human (SH-SY5Y) immortalized neuronal cell lines, H584R and FYHR exhibited increased transduction relative to AAV9 (Fig. 3E and 3F). This phenotype was particularly pronounced in HEK293 cells, which are historically less permissive to AAV9, where H584R and FYHR demonstrated significantly increased transduction compared with wild-type AAV9 (Figure S3). Together, these findings suggest that H584R confers a generalized enhancement in transduction that is not restricted to a single cell type.

### H584R alters AAV capsid specific cell-surface binding behavior

To directly determine whether the mutations altered cell-surface interactions, we performed a capsid binding assay across increasing vector doses in CHO lec2 cells. At lower doses, binding of the variants was broadly comparable to AAV9; however, at doses exceeding 1 × 10^5^ vg/cell, H584R and FYHR demonstrated increased particle binding that continued to accumulate over time, producing binding kinetics distinct from wild-type AAV9 (Figure S4). These findings indicate that H584R alters the interaction of the capsid with the cell surface and may increase engagement of one or more additional binding partners.

### Enhanced transduction is not explained by HSPG dependence

Acquisition of positively charged capsid residues can promote heparan sulfate proteoglycan (HSPG) interactions, therefore we next tested whether the H584R substitution increased transduction through HSPG-mediated uptake. HEK293 cells were transduced in the presence or absence of soluble heparin competition. Heparin treatment produced only a modest reduction in transduction of AAV9, F535Y, H584R, and FYHR, whereas transduction by AAV2, which relies strongly on HSPG binding, was nearly completely inhibited (Figure S5.). Thus, the increased activity of H584R and FYHR is unlikely to result from acquisition of substantial HSPG-dependent entry. Collectively, these findings indicate that the variants retain sensitivity to AAV9-associated galactose availability but that neither altered galactose utilization nor acquisition of HSPG dependence is sufficient to explain their enhanced transduction. This raised the possibility that H584R alters interactions with additional, unidentified cell-surface factors.

### FYHR exhibits biophysical properties distinct from either single mutant

Although altered cell-surface binding could contribute to the increased transduction observed with H584R and FYHR, these changes do not exclude additional impacts on capsid stability, intracellular trafficking, or uncoating. To explore these features further, we used differential scanning fluorimetry (DSF) to determine whether the mutations altered capsid thermal stability under conditions relevant to AAV characterization and intracellular trafficking^39,40^. At physiological pH (pH 7), FYHR exhibited a modest reduction in thermal stability compared with AAV9, whereas neither F535Y nor H584R alone produced a significant thermal shift (Figure S6A & S6B). Similarly, particle charge measurements revealed a significant shift toward a more negative surface charge for FYHR that was not observed with either single mutant (Figure S6C).

Together, these findings indicate that the combined F535Y and H584R substitutions produce biophysical properties that are not recapitulated by either mutation independently. Because AAV particles encounter progressive acidification during endosomal trafficking, we next examined capsid thermal stability across decreasing pH conditions. AAV9 maintained relatively consistent thermal stability from physiological to moderately acidic conditions before exhibiting a pronounced reduction at pH 4 (Figure S6B). FYHR displayed a distinct response, with modestly increased thermal stability at pH 6 and pH 5 followed by a similarly pronounced reduction at pH 4 (Figure S6B). These results suggest that the combination of F535Y and H584R alters the response of the capsid to environmental acidification and may influence conformational transitions encountered during intracellular trafficking.

Together, these experiments reveal distinct but potentially complementary contributions of the two FYHR mutations. H584R is sufficient to drive much of the enhanced transduction phenotype and alters capsid-cell binding without conferring substantial HSPG dependence or disrupting responsiveness to terminal galactose. In contrast, measurable changes in capsid thermal stability, pH-dependent behavior, and particle charge emerge only when F535Y and H584R are combined. Thus, FYHR appears to couple an H584R-driven enhancement in cellular interaction and transduction with emergent biophysical properties produced by the combined mutations, providing a potential mechanistic basis for its improved gene-delivery phenotype.

### FYHR has increased gene delivery to the CNS of mice following intrathecal administration

Traditional capsid engineering validation strategies are notoriously laborious, time intensive, and expensive. To accelerate and alleviate the cost and time burden that is required to directly compare multiple vectors, we developed a multiplexed reporter vector system that allows the direct comparison of five capsids within a single administration in the same animal. The system comprises of six main elements: (a) self-complementary ITR elements (one wildtype ITR and one mutated ITR); (b) a moderate ubiquitous promoter (UsP); (c) an N-terminal unique 3X protein epitope tag (FLAG, V5, HA, HIS, or MYC) linked to an (d) enhanced green fluorescent protein (eGFP); (e) a unique 400 bp 3’ UTR barcode; and (f) a small synthetic Poly-A element. Therefore, we decided to package our benchmark capsid, AAV9, with the FLAG-ITGBC1, FYHR with V5-ITGBC2, HRLI with HA-ITGBC3, a control capsid variant (“NSKN”) with HIS-ITGBC4, and H584R single mutant variant with MYC-ITGBC5. The pooled reporter vectors were administered as a single bolus intrathecal lumbar cistern injection of 1 x 10^11^ vg/capsid (5 x 10^11^ vg total) in adult mice (Fig. 4A). There was a significant increase in vector genomes delivered by FYHR across CNS tissues, but surprisingly this was not seen with the single mutant H584R when compared to AAV9 (Fig. 4B). All peripheral tissues, except liver, showed no significant differences in vector genome delivery compared to AAV9 (Fig. 4C). The liver showed 3-fold (FYHR) and 2-fold (HRLI and H584R) higher vector genome delivery when compared with AAV9 (Fig. 4C).

**Figure 4.**
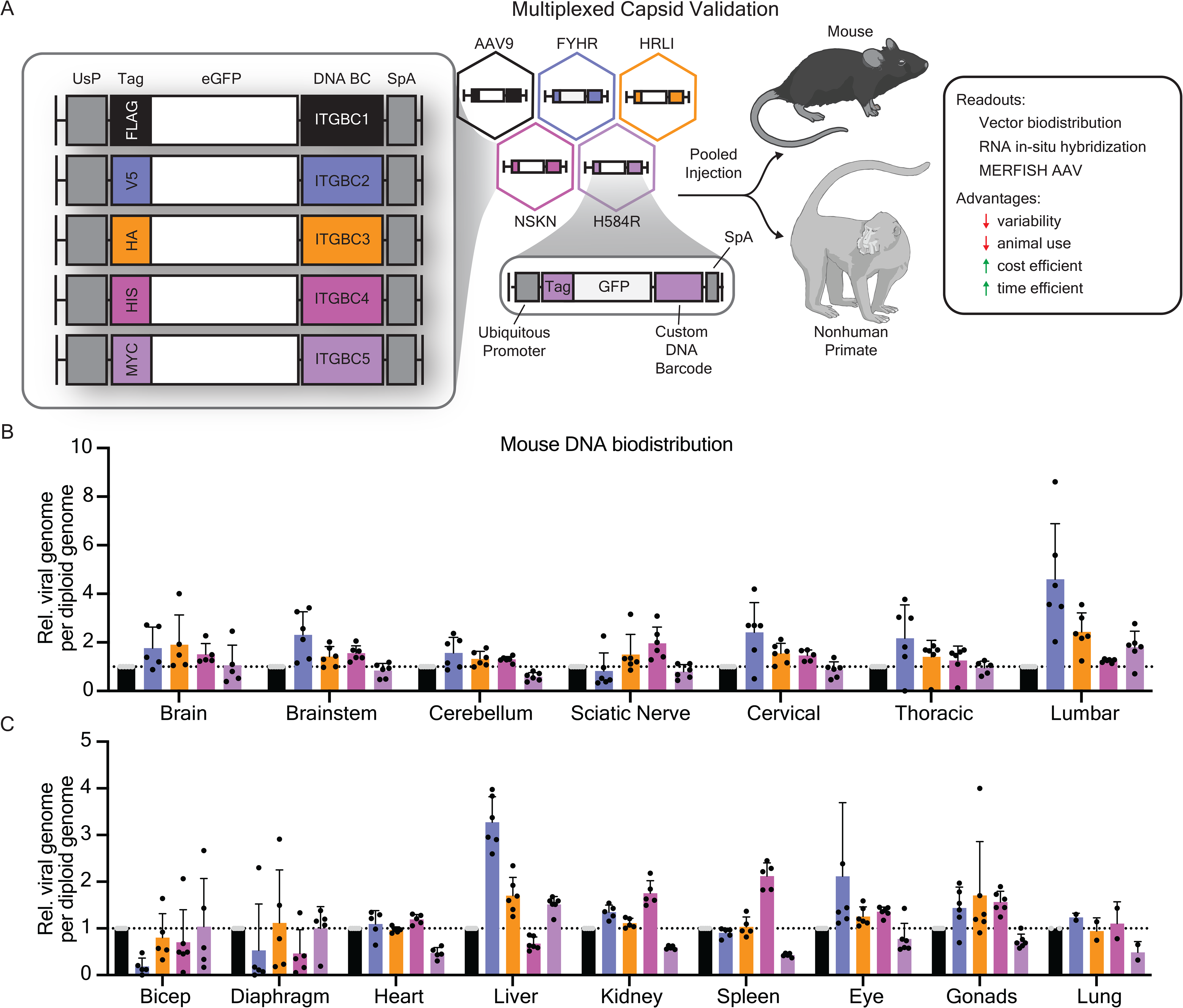
Vector DNA biodistribution of capsid variants in mice. (A) Schematic of the multiplexed capsid validation reporter system. A self-complementary ITR configuration flanks a moderate ubiquitous promoter (UsP) expressing an epitope tagged eGFP with a 3’UTR unique DNA barcode (ITGBC1-5) and a small synthetic PolyA signal. Five capsids are injected as a pool into mice and nonhuman primates. In wildtype C57BL/6J mice (n = 6), the capsid pool was administered by a lumbar intrathecal route (see methods). (B-C) Biodistribution reported as viral genomes per diploid host genome relative to AAV9 for CNS (B) and peripheral tissues (C). Dashed line = 1 (normalized AAV9 of each mouse).

Some limitations of the reporter multiplex system were identified that should be considered. Protein expression was not similar across all reporter constructs, such that the V5 and HA tagged vectors had significantly lower GFP expression compared to FLAG, HIS, and MYC (Figure S7 & S8). This rendered expression comparisons at the protein level uninformative to evaluate FYHR (V5). RNA expression was similar across constructs, except that ITGBC4 (NSKN) had slightly lower transcript expression than the rest (Figure S7 & S8).

### Increased FYHR gene expression correlates with vector genome delivery in mice

To assess vector-mediated transgene expression, we utilized *in situ* hybridization with RNAScope probed against the barcodes transcriptionally expressed in the 3’ UTR of each reporter. Consistent with the increased vector genome delivery, RNAScope revealed FYHR-mediated transgene expression enhancement in the brain when compared to AAV9 (Fig. 5A). Image quantification confirmed comparable enhancement of gene expression for FYHR as we observed in vector genome delivery within several regions of the brain and spinal cord, but expression wasn’t significantly increased for the other variants (Fig. 5B-G). While RNAScope provided broad gene expression information, we sought to determine if the tropism of FYHR had significant differences in cell type transduction within the brain. Multiplexed fluorescence in-situ hybridization (MERFISH) was used to evaluate the single cell tropism of FYHR with respect to AAV9. We designed a custom 140 bp gene panel that included all 5 of our custom barcodes and cell type identification markers (see Methods). We confirmed that we could identify major cell types in the sagittal mouse brain via this method (Fig. 5H & 5I). Overall, we found that FYHR had very similar cell type targeting to AAV9, except endothelial cells were transduced less by FYHR compared to AAV9, which seemed to re-distribute to a higher proportion of astrocytes (Fig. 5J). Spatial visualizations of the five capsids revealed no significant differences in terms of subregional brain targeting (Figure S9).

**Figure 5.**
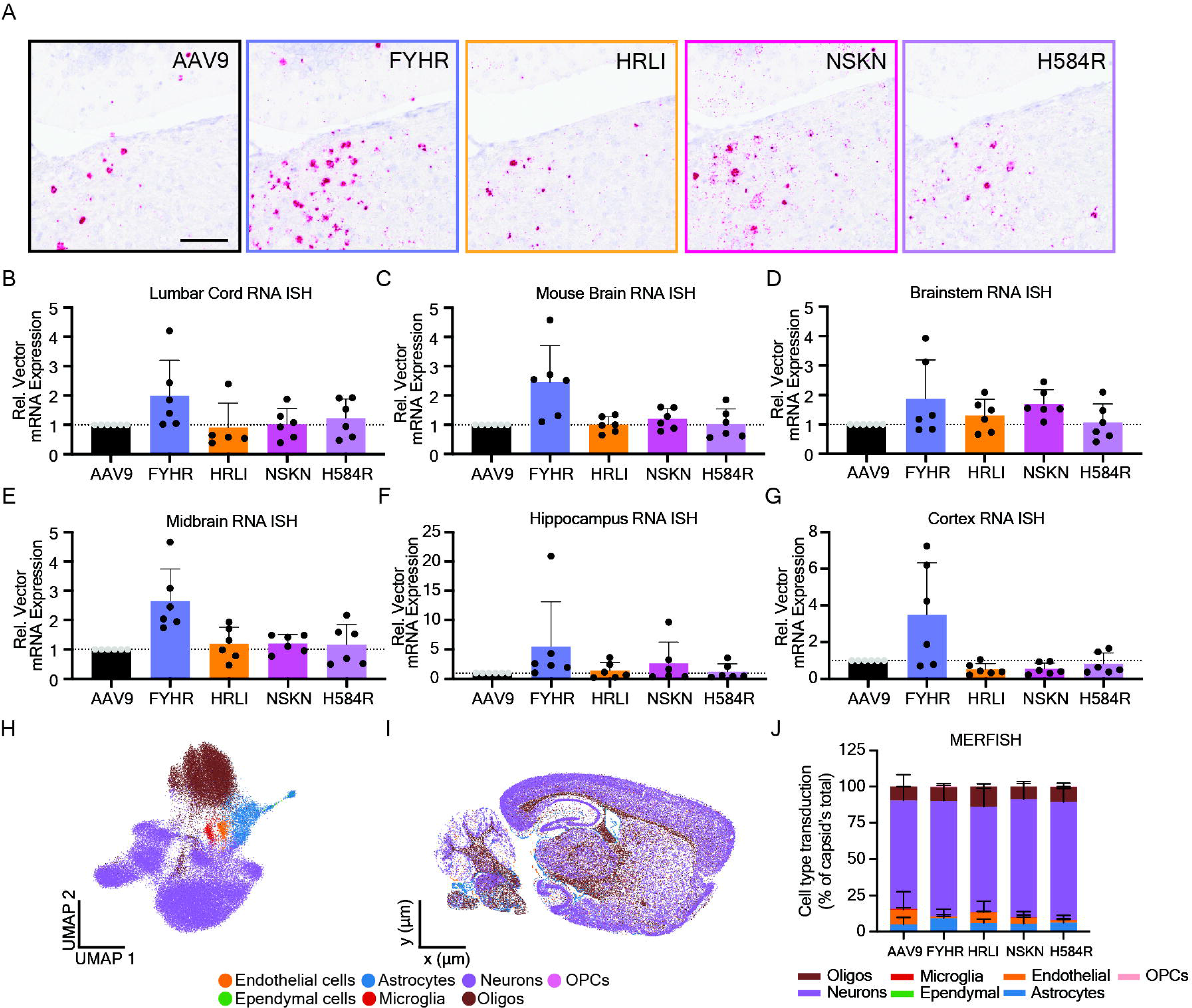
Vector RNA expression and single cell transduction profile of AAV9 variants in mice. (A) Representative images of RNA in situ hybridization (RNAScope) probed against 3’ UTR barcodes for each capsid in serial 5 µm sections of the mouse hindbrain (sagittal). Blue = nuclei; pink = AAV transgene RNA. (B-G) RNAScope image quantification of lumbar cord (B), whole brain (C), brainstem (D), midbrain (E), hippocampus (F), and cortex (G) (n=6 mice). (H) Unifold manifold approximation (UMAP) of MERFISH data with cells colored by cell subclass identification. (I) Spatial maps of the sagittal mouse brain with cells colored by cell subclass identification. (J) Histoplot displays cell type transduction percentage of total cells transduced by each capsid. (n=2); Scale bar = 100 µm.

### FYHR demonstrates improved CNS gene delivery in nonhuman primates

Next, we investigated whether FYHR would improve gene delivery in nonhuman primates. Two rhesus macaques of similar age were infused with the same pool as described in mice at 3 x 10^13^ vg per capsid (1.5 x 10^14^ vg total), with the exception that one NHP was injected with FYHR at half the dose of AAV9 (Methods). Vector distribution across 8 brain samples showed increased gene delivery of 2- to 4-fold for FYHR, HRLI, and H584R variants compared to AAV9 (Fig. 6A). Three separate samples of the cerebellum had 4-fold improved gene delivery and every region of the spinal cord showed closer to 5-fold improvement over AAV9 (Fig. 6A). As in the mouse validation experiments, peripheral tissues showed no difference in gene delivery compared to AAV9, except the liver which showed a 2-fold increased gene delivery (Fig. 6B). Finally, we found that DRGs, lumbar roots, ulnar nerve, and femoral nerves showed 2- to 6-fold increased vector genomes of FYHR compared to AAV9 (Fig. 6C). When correcting for the half dose of FYHR in the one NHP, both animals showed consistent fold differences of the capsid variants compared to AAV9.

**Figure 6.**
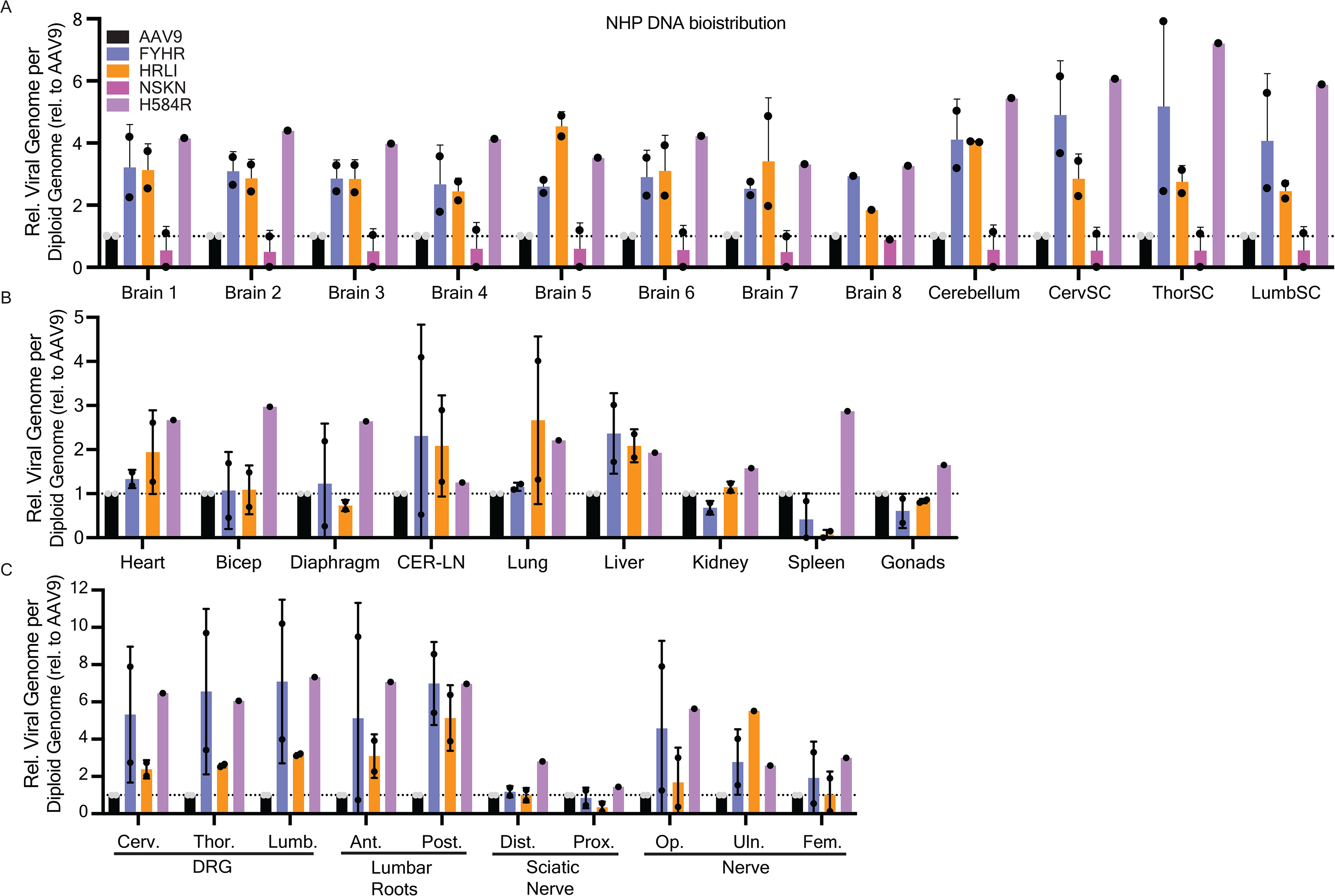
Vector genome biodistribution of AAV9 capsid variants in nonhuman primates. In adult rhesus macaques (n = 2), the capsid pool was administered by a lumbar intrathecal route (see methods). (A-C) Biodistribution is reported as viral genomes per diploid host genome relative to AAV9 for CNS (A), peripheral tissues (B), and PNS (C). Dashed line = 1 (normalized AAV9 of each NHP).

### FYHR delivered gene expression correlates with vector genome delivery in nonhuman primates

NHP RNAscope of brain samples similarly revealed increased transgene expression mediated by FYHR over AAV9 (Fig. 7A). Quantification of the RNAscope expression showed similar enhancement as the vector genome analysis across all brain and spinal cord regions of approximately 4- to 6-fold improvements for FYHR compared to AAV9, along with 3-fold increased liver expression (Figure S10). Notably, one of the two NHPs showed substantially higher vector genomes across DRGs and lumbar roots (approximately 8-11 times higher than AAV9). However, RNAscope analysis of this NHP’s DRG showed approximately 3 times higher expression with FYHR compared to AAV9, consistent with increases across the rest of the CNS (Figure S10. Subsequently, we designed a second custom 140 gene panel for the NHP brain to characterize the single cell tropism of our engineered capsids alongside AAV9. We identified all major cell types in the NHP brain within the cortex (Fig. 7E, 7F & Methods). Upon quantification of capsid transduced cell types, we found that FYHR had a comparable transduction profile to AAV9. Similar to the results in mice, we saw that the proportion of transduced endothelial cells was reduced in FYHR when compared with AAV9 (Fig. 7G).

**Figure 7.**
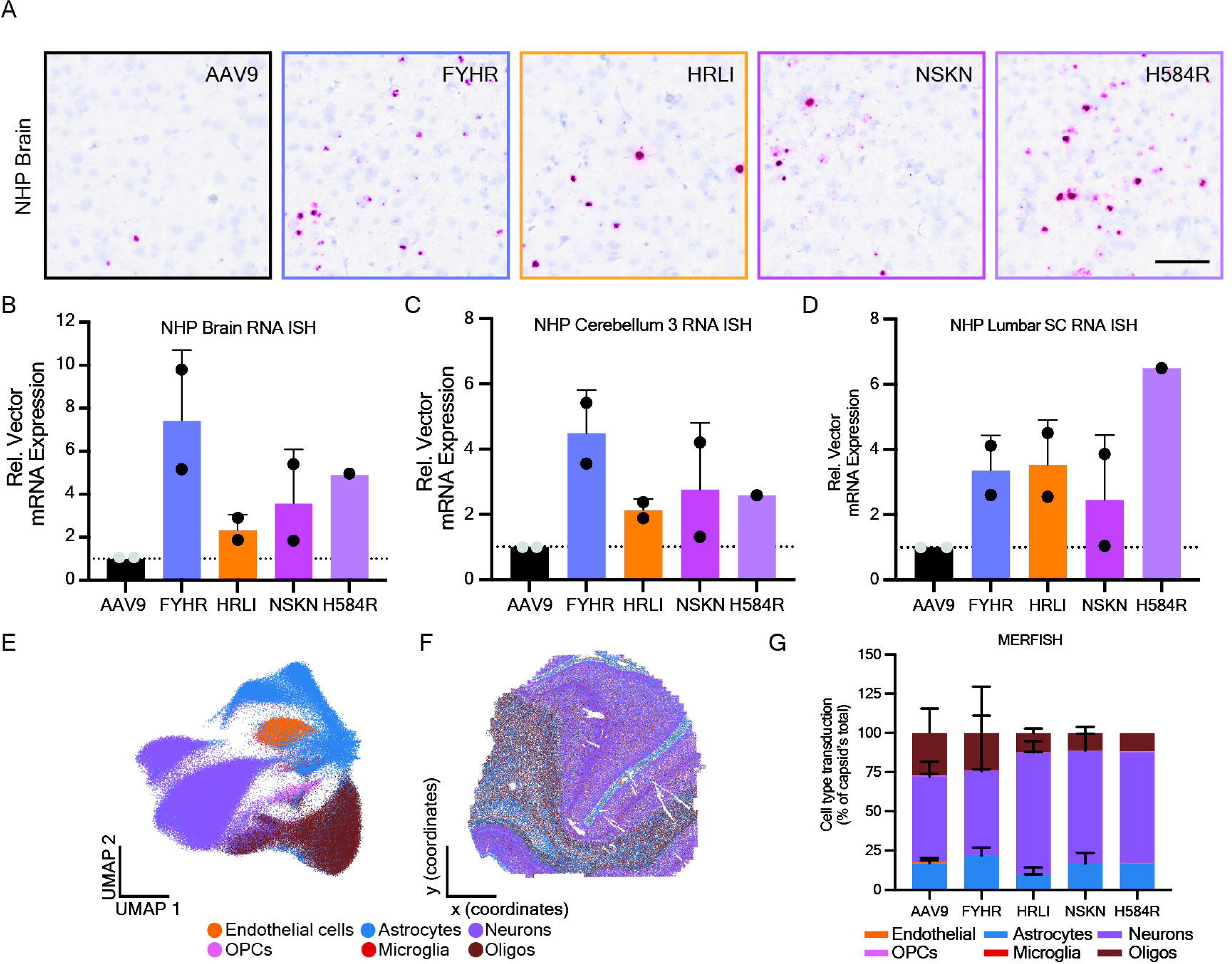
Vector transgene expression and single cell transduction profile of AAV9 capsid variants in nonhuman primates. (A) Representative images of RNA in situ hybridization (RNAScope) probed against 3’ UTR barcode for each capsid in serial 5 µm sections of the NHP cortex (sagittal). Blue = nuclei; pink = AAV RNA. (B-G) Aggregated RNAScope image quantification of 4 brain regions (B), 3 cerebellum regions (C), and 5 whole lumbar cord cross-section (D), (n=2 NHP). (E) Unifold manifold approximation (UMAP) of MERFISH data with cells colored by cell subclass identification. (F) Spatial maps of the NHP brain section with cells colored by cell subclass identification. (G) Histoplot displays cell type transduction percentage of total cells transduced by each capsid. (n=2); Scale bar = 400 µm.

## DISCUSSION

AAV mediated gene therapies continue to hold promise for the treatment of genetic diseases. While AAV9 has been the dominant capsid serotype for CNS delivery^1^, it continues to show limitations in its ability to transduce a large percentage of cells across the entire CNS. As such, others have proceeded to engineer AAV9 for enhanced delivery to several different tissues, including the CNS^28,29,31,41^.

Most engineering strategies have focused on small peptide insertion sequences into amenable surface exposed loops with the goal of engaging a new receptor or cell surface attachment factor in order to increase the efficacy of gene delivery^28,41–43^. However, AAV’s transduction pathway involves additional critical steps beyond attachment to the cell surface, which can potentially be leveraged to increase the potency of vectors. While promising, peptide insertion libraries affect isolated regions of the capsid that may limit the degree of evolution that can occur across the complex multi-subunit 3-dimensional structure of the capsid. However, strategies that explore manipulations across larger stretches of VP3, such as DNA shuffling and random mutagenesis, have historically been challenging and limited by sequencing technology read length limitations and throughput bottlenecks. To address some of these challenges and limitations we designed an AAV9-based library that sought to explore evolutionary pressures across the larger sequence space of the AAV9 VP3 protein. To increase the pool of functional variants in the library, highly conserved structural domains were avoided as well as variable regions that span AAP or are necessary in genome packaging (VRI and VRII)^37,44,45^. Furthermore, we further demonstrate the utility of single molecule real-time sequencing (SMRT seq) for the identification of AAVs from higher complexity libraries previously generated through capsid shuffling^35^, and presently, through controlled randomization.

Viral tropisms beyond tissue targeting have posed a challenge in capsid engineering, specifically with cross-species compatibility^46^. Others have previously tackled this by sequentially screening and applying directed evolution through multiple species such as screening through pigs then filtering through mice, and finally evaluating those recovered variants through nonhuman primates^47^. Each round of selection increases the time it takes to identify a variant of interest. To reduce time, we performed screens in parallel across mice and nonhuman primates. The lead candidate from these screens, AAV9fyhr, was likewise identified in a single round of selection (Fig. 1) and confirmed in a second round through human iPSC differentiated cortical neurons (Fig. 2). Through our screens, we identified hotspots that may be more critically important than others for brain transduction of mice and nonhuman primates (Figure S11). Future work aims to explore the incorporation of other identified amino acid variants identified in the screens, to rationally design future iterations of FYHR.

FYHR demonstrates that minimal modifications can be sufficient to increase vector transduction. We show that a single amino acid change H584R is sufficient to increase binding at the cell surface (Figure S4.) and ultimately increase gene expression in vitro and in vivo. Although F535Y did not directly mediate changes in transduction, biophysical properties such as computational predictions, viral stability, and zeta potential measurements reveal the necessity of F535Y (Figure S1, S6, & S12) that are inexplicable by structural reconstructions using Cryo-EM. Initially, Alphafold predicted a specific rotamer confirmational change at the F535Y position. We tested the hypothesis that F535Y produces a detectable rearrangement of the VP3 fold or of the inter-subunit interfaces by comparing the 1.87 Å FYHR reconstruction to the 2.35 Å AAV9 wild-type reconstruction (PDB: 7MT0^48^) determined in parallel. Cα superposition of the refined chain-A models yielded a global RMSD of 0.363 Å (513 pruned pairs, residues 219–736; 0.425 Å across all 516 pairs). This outcome is consistent with published AAV2 point-mutant capsid structures L336C (PDB 6E9D^49^), R432A (PDB 5IPK^50^), and R404A (PDB 8FZN^51^), which all retain wild-type capsid architecture with no significant Cα deviations despite producing substantial functional phenotypes. It is noteworthy that the Alphafold-predicted conformation of F535Y was not consistent with the cryo-EM structure, perhaps as a cautionary note to not overly rely on current *in silico* prediction models. Overall, we could not conclude any impact of F535Y to the overall capsid structure.

Importantly, these two mutations did not disrupt the binding of n-terminal galactose, but they were sufficient to engage HSPGs at a lower level compared to AAV2 (Figure S5.). Furthermore, this dual glycan binding may be contributing to part of its enhanced transduction, but the transduction is not exclusively dependent on HSPG (Figure S5). Overall, our studies did not identify (or exclude) a novel receptor for FHYR, but future studies could explore the possibility of additional receptor, cell surface binding partner(s), or additional intracellular host factors facilitating an increase in transduction.

Traditional methods of capsid validation remain laborious, resource heavy, and time intensive. To develop an accelerated validation platform, we developed a first-generation multiplexed reporter vector system that enabled head-to-head comparisons of up to 5 vector designs within the same animal. Additionally, the validation and characterization of novel engineered capsids pose the question whether the same tissues or even cell types are transduced. Implementing long DNA barcodes enabled the use of single cell spatial transcriptomics, RNA *in situ* hybridization (RNAScope), and DNA biodistribution. One drawback of this first-generation system is that the protein expression was technically challenging to detect in vivo due to its weaker expression level, and two of the designs (V5 and FLAG) resulted in lower GFP protein expression. Four out of the five had comparable RNA expression, with the BC4 vector showing significantly reduced (∼half) RNA expression (Figure S7 & S8). Optimization of these reporters to have fully matched RNA and protein expression levels, under the control of a stronger promoter, will increase their utility.

Lastly, we identified an engineered capsid variant that demonstrated increased CNS transduction in both mice and nonhuman primates after lumbar intrathecal administration. Furthermore, the mutations enriched *in-vivo* were found enriched in iPSC derived human cortical neurons, providing a strong rational for clinical development. While the liver was not de-targeted, the remaining peripheral tissues showed comparable transduction to AAV9. Single cell transcriptomics shows similar proportions of cell types transduced as AAV9, which further corroborates our in vitro findings that AAV9fyhr has increased transduction potency without large changes to its overall tropism. One difference from cell types transduced in the brain was the population of endothelial cells and astrocytes. FYHR transduced lower proportions of endothelial cells, and effectively replaced them with an increased number of transduced astrocytes. We don’t have an explanation for this change, but we might speculate that this could be from vector circulating in the blood after the intraCSF injection, which could target astrocytes at the BBB if the endothelial cells aren’t being transduced. Indeed, we observed that FYHR circulates in the blood at higher amounts within the initial ∼2 hours following the intrathecal injection in mice (Figure S13).

In summary, our study presents a minimally modified AAV9 variant that is compatible in both mice and nonhuman primates with enhanced transduction of the CNS after lumbar intrathecal administration. Our results build on structural and functional understanding of the AAV9 capsid toward developing whole capsid engineered variants. We demonstrate that two amino acids are sufficient to have pleiotropic effects that span biophysical properties of the viral particle, cell surface lectin binding, gene delivery, and tissue tropism. Furthermore, we demonstrate a multiplexed approach to capsid validation that incorporates single cell transcriptomics to characterize cell type targeting at a more granular resolution. Overall AAV9fyhr displays a similar CNS tropism compared to AAV9, but with a transduction potency that is ∼2-3 fold higher in mice and ∼4-6 fold higher in nonhuman primates. Based on these properties, we suggest that AAV9fyhr can be adapted into current preclinical developmental paradigms of intrathecal gene therapy approaches, leveraging prior AAV9 data and using AAV9fyhr to either lower the effective dose or achieve higher transduction efficiencies at the same AAV9 dose.

## METHODS

### Plasmids

The pSSV9 vector, previously described^35^, contains 2 main elements flanked by WT AAV2 ITRs: (i) an AAV2 Rep gene; and (ii) WT AAV Cap gene (in this case using AAV9 instead of AAV2). The multiplexed reporter plasmids generated for pooled capsid validation studies contain five main elements flanked by an upstream mutated AAV ITR and downstream WT AAV ITR, driving packaging of self-complementary genomes: (i) a ubiquitous promoter, also known as “UsP”^52^; (ii) a 3X protein epitope tag [FLAG, V5, HA, His, or Myc] with a C-term GS linker tagging; (iii) an enhanced GFP or “eGFP” ; (iv) a 3’ UTR 400 bp barcode, “ITGBC1-5”, designed in-silico to have no homology to mouse, rat, rhesus macaque, cynomologous macaque, canine, or human; and a (v) Synthetic PolyA, also known as “SpA”^53^.

### Capsid library generation

The AAV9 controlled random mutagenesis library was designed by subjecting the wildtype AAV9 sequence to random mutagenesis but limited to only variable regions III-IX. Synthesis of the controlled mutagenesis was performed by Thermo Fisher’s GeneArt services, targeting approximately ∼3-5 mutations per clone. The library was generated with a target >1 x 10^11^ independent molecules of diversification and cloned into the pSSV9 plasmid backbone by HindIII and XbaI traditional restriction enzyme cloning as previously described^35^.

### Animals

All procedures were performed in accordance with protocols approved by the Institutional Animal Care and Use Committee of the University of Texas Southwestern Medical Center (UT Southwestern Medical Center), an AAALAC-accredited facility. C57BL/6 breeders were obtained from Jackson Laboratories and maintained at UT Southwestern Medical Center. Animals were maintained under controlled environmental conditions, 12-hr light-dark cycles, and provided with food and water ad-libitum. Rhesus macaque studies were conducted at the Tulane National Biomedical Research Center (TNBRC), which is fully AAALAC-accredited, with approval by the Tulane University Institutional Animal Care and Use Committee (IACUC) and the Tulane University Institutional Biosafety Committee (IBC).

### Viral vector production

All AAV productions utilized Expi293F cells (Thermo; cat#: A14527) with Expi293 growth medium (Thermo) by traditional triple transfection using FectoViR-AAV (Polyplus) and purified by CsCl ultracentrifugation, adapted with slight modification from a previously described method^54^. The AAV9 controlled random mutagenesis (CRM) library was produced in two rounds of production: (i) Triple transfections using Ad helper pALD-X80 (Aldevron), AAV2 helper, and AAV9 CRM library in a 1:10:1 (respectively), such that the library is preferentially packaged into an AAV2 capsid, after purification, also referred to as “shuttle” library; (ii) the “shuttle” library was then used in a consecutive round of production using WT Ad5 (Baylor College of Medicine viral vector core) at a low MOI such that a single AAV shuttle particle transduced a single cell in order to avoid AAV mosaic assembled particles, after purification, also referred to as “final” library. Recombinant AAV capsid variants were produced as previously described^54^ using Ad helper pALD-X80 (Aldevron), AAV Helper, and ITR containing reporter plasmids.

Clarified lysates were subjected to one round of step gradients followed by two rounds of continuous CsCl ultracentrifugation. Peak fractions containing full encapsidated particles were identified by qPCR and dialyzed with a modified PBS (1X PBS, 5% sorbitol, 0.001% F68 pluronic).

### Capsid library screening and capsid variant recovery

All rhesus macaques were screened for pre-existing AAV9 neutralizing antibodies prior to capsid library injections, and only seronegative animals were used. NHPs received a 1.5 mL infusion of the capsid library via lumbar intrathecal (IT) administration at 4 x 10^13^ vg per animal. C57BL/6 received a single bolus 5µL IT injection at 1 x 10^11^ vg per animal. Animals were euthanized 2 weeks after capsid library administration and perfused with 1X phosphate buffered saline. After perfusion, tissues were frozen and stored at -80 °C for long term storage. Nuclei was isolated from frozen tissue using Nuclei EZ Prep (Sigma-Aldrich) kit. DNA from nuclei isolations was purified by DNeasy blood and tissue kit (Qiagen). PCR amplification of capsid sequences was done using CloneAmp Hifi PCR master mix (Takara).

### Single molecule real-time (SMRT) sequencing

PCR amplicons of full length capsid sequences (2.2 kb cap gene) were used to generate SMRT sequencing libraries using the SMRTbell prep kit (PacBio) via manufacturer’s specifications. SMRT seq libraries were sequenced on a Sequel IIe sequencer at UT Southwestern Medical Center’s NGS sequencing core facilities. Data obtained from the PacBio sequencer was checked for quality using SMRT Link. The subread bam files produced by SMRT Link were converted to Circular Consensus Sequences (CCS) FASTA files using ccs (v3.3.0) from PacBio tools. The CCS FASTA files were then collapsed and counted using a python script. Unique DNA sequences were then queried for potential ORFs and translated to protein sequences using getorf tool (getorf -minsize 2100 -find 1 -table 0) from EMBOSS. Protein sequences were further collapsed and counted to obtain the final protein frequencies (Counts/Total).

### Site-directed mutagenesis

For single variant characterization, we performed site-directed mutagenesis to AAV helper plasmid (pAAV9) using Quikchange II XL site-directed mutagenesis kit (Agilent) per manufacturer’s specifications. Primer design was done using Agilent’s QuikChange Primer Design Tool (https://www.agilent.com/store/primerDesignProgram.jsp).

### Capsid validation in vivo

Pooled AAV lumbar intrathecal injections were performed in adult (6-8week old) C57BL/6J mice (n=6; 3M;3F) at 5 x 10^11^ vg total (1 x 10^11^ per capsid). Mice were euthanized 4 weeks post-injection. Half of tissues collected were flash frozen for biodistribution and the other half were fixed in 10% neutral buffered formalin, 24 hours post fixation, moved to 70% ethanol and processed for paraffin embedding. Pooled AAV lumbar intrathecal injections were performed in juvenile rhesus macaques (n=2; 1M;1F) 12-13 years of age. In the male NHP (12.38 years of age) 1.5 mL of capsid pool (3 x 10^13^ vg per capsid for AAV9, HRLI, NSKN, and H584R; 1.5 x 10^13^ vg of FYHR) was administered. For the female NHP (12.83 years of age), 1.5 mL of capsid pool (3 x 10^13^ vg per capsid for AAV9, HRLI, NSKN, and FYHR) was administered. Tissues were collected and frozen for vector biodistribution and Tissues were fixed in 10% neutral buffered formalin. After fixation, tissues were moved to 70% ethanol and processed for paraffin embedding.

### RNAScope

Serial sections for RNA in situ hybridization were cut at 5 μm thickness. Each tissue group for staining and capsid comparison was collected within a 50 μm range. RNAScope (ACDBio) was used to stain for the capsid barcode using custom RNAScope probes following manufacturers specifications. Hematoxylin was used as a counter stain for nuclei. Images were collected using a Hamamatsu Nanozoomer S60 digital slide scanner at 60X. Images were visualized and prepared for publication using NDPview2 software. Quantification of RNAscope was performed using HALO (Indica labs) pixel quantification software. Each section was quantified for hematoxylin and RNA staining pixel quantification. RNA staining was normalized to hematoxylin for every tissue. Subsequently, each tissue was then normalized and compared relative to the AAV9 condition for each tissue per animal.

### Vector biodistribution

DNA was extracted with DNeasy blood and tissue kit (Qiagen). Viral vector genomes were quantified by qPCR using a LightCycler 480 Instrument II (Roche) probing for the DNA barcodes. Mouse or rhesus Gapdh was probed and used as a normalizing gene to determine viral genome per diploid genome copy.

Cryo-electron microscopy sample processing, data collection, and structure determination Concentrated AAV (4.12 x 10^13^ vp/mL) was diluted ¼ then 3.5 µL of diluted AAV was applied to glow-discharged carbon grids (Quantfoil Cu R1.2/1.3, 300-mesh). Grids were plunge-frozen in liquid ethane using an FEI Mark IV Vitrobot (Thermo Fisher Scientific) (4°C, 100% humidity, 45 second delay, 4 second blot time, blot force 0). Grids we clipped and screened on a Talos Arctica (FEI). The grids were then loaded onto a 300 kV Titan Krios (FEI) micrscope for imaging. Movies consisting of 58 frames were acquired using a Gatan K3 summit detector. The total dose administered was 58 e^-^/ Å^2^, and the super resolution pixel size was 0.415 Å. To eliminate inelastically scattered electrons, a Bioquantum energy filter with a slit width of 20 e^-^V was employed. Movies were binned by two, motion corrected, and dose weighted using the motion correction implementation provided by RELION 4.0^55^. Aligned and dose-weighted micrographs were then used to estimate the contrast transfer function (CTF) using CTFFIND-4.1^56^. Four micrographs were manually picked, yielding 195 particles used to train a neural network, TOPAZ^57^. The trained model was used to pick the full dataset of 5,010 micrographs with a Topaz threshold value of 5. Autopicked particles were extracted using a box size of 380 pixels (∼317 Å), without rescaling. Two rounds of reference-free 2D classification were performed using a mask diameter of 300 Å. The highest resolution class averages were selected, yielding 224,811 particles for 3D refinement. For 3D refinement, the published AAV9 map (EMDB-23973, PDB 7MT0) was low-pass filtered to 60 Å using relion_image_handler and used as the initial reference. A first 3D auto-refinement was conducted with C1-symmetry. A soft-edged solvent-flattened mask was generated from the first reconstruction, and a second 3D auto-refinement was performed with solvent flattening. I1 icosahedral symmetry was imposed in the third 3D auto-refinement. Bayesian polishing was performed, and the polished particles were used for a fourth round of refinement. Three consecutive rounds of CTF refinement were performed: (1) anisotropic magnification, (2) per-particle defocus and astigmatism, and (3) higher-order aberrations. A fifth refinement was performed with the CTF-corrected particles. Post-processing with automated B-factor estimation (-65.0 Å) and solvent-corrected FSC yielded a final resolution of 1.87 Å at the FSC = 0.143 with 224,811 particles. For model building and refinement, the published AAV9 structure (PDB: 7MT0)^44^ was used as the starting model. Substitutions F535Y and H584R were introduced using ChimeraX^58–60^. The single-chain VP3 monomer (residues 219-736) was co-registered to the cryo-EM map using ChimeraX, then subjected to Phenix real-space refinement^61,62^. Model validation was performed using MolProbity^63^. The refined single-chain model was expanded to the full 60-subunit capsid using I1 icosahedral symmetry operators in ChimeraX. The final map-model correlation coefficient for the chain A model was 0.892.

### MERFISH

Two custom 140 gene panels were designed to discriminate major cell types in the adult mouse and nonhuman primate brains as well as our barcoded AAV multiplex reporter designs (ITGBC1-5). Tissues were fixed in 10% neutral buffered formalin for 24 hours at room temperature. Tissues were then processed for paraffin embedding. Paraffin blocks were sectioned at 5 µm thick sections onto MERSCOPE slides and allowed to dry overnight. Tissues were processed and imaged per manufacturer’s specifications. After imaging, datasets were re-segmented using custom trained cell segmentation models for mouse brain or nonhuman primate brain tissue in Cellpose 2^64^. High quality MERFISH data filtering was done by using cells with >10 transcripts per cell. Analysis and visualization of MERFISH data was done using Squidpy^65^, Anndata^66^, and custom python scripts.

### Viral Transduction Assays

Cells (HEK293T, Lec2, SH-SY5Y, Neuro-N2A, Cos7, and iPSC-derived cortical neurons) were seeded into 96-well flat bottom plates with 10,000 cells per well. 24 hours after seeding, cells were treated with 1 x 10^4^, 1 x 10^5^, or 1 x 10^6^ vg per cell. Cells were collected 48 hours after virus treatment for immortalized cells, or seven days post transduction for iPSC derived cortical neurons. In brief, growth medium was removed, cells were gently washed with 1X phosphate buffered saline, dissociated with 0.25% trypsin, neutralized with outgrowth medium, and then transferred to a round bottom 96-well plate before pelleting by centrifugation. Medium was removed and cells were fixed with 4% PFA at room temperature for 15 minutes. PFA was removed and cells were resuspended using 1X PBS. Cells were quantified for GFP+ cells using an Attune Cytpix (Thermo) flow cytometer.

### Manufacturability assays

Adapted from previously described^54^, Expi293F cells were seeded at 1 x 10^6^ cells/mL in 30 mL shaker flasks. 24 hours after cell seeding, triple transfections using Ad helper pALD-X80 (Aldevron), AAV helper, and ITR plasmid (pTRS-ks-Cbh-eGFP-BGHpA) at a 1:1:1 molar ratio with FectoVir-AAV (Polyplus) 1uL:1ug of DNA for every 1 x 10^6^ cells. 1 mL sample of culture was collected at 48-hours post transfection. Samples were subjected to three freeze/thaw cycles and vortexed after each thaw. After the final thaw, cell lysate was clarified by centrifugation. 10 uL of each sample was DNase I treated for 2 hours at 37 °C, quenched with 6 uL of 0.5 M EDTA (Sigma), heat inactivated for 10 mins at 75 °C, and then treated with proteinase K. Samples are then brought up to 1 mL with DNase/RNase free Ultrapure water (Invitrogen) and further diluted 1:200 in DNase/RNase free Ultrapure water (Invitrogen). Viral genomes are measured using a linearized plasmid standard and absolute quantitative PCR with PowerUp 2X SYBR Green master mix (Applied Biosystems) on an QuantStudio 5 real-time qPCR machine (Applied Biosystems).

## Statistical Analysis software

All statistical analyses were performed in GraphPad Prism (version 11.0.0). For all statistical analysis, significance is represented as *p<0.05, **p<0.01, ***p<0.001, ****p<0.0001; non-significance p>0.05.

## Data availability statement

Source data for each relevant figure is provided in a source data file. Additional data that support the findings of this study are available upon reasonable request from the corresponding author, Steven J. Gray.

## Supporting information

Supplemental Methods and Supplemental References

## Acknowledgments

We would like to thank the members of the UT Southwestern Medical Center’s Next Generation Sequencing Core for the SMRT sequencing library preparation and sequencing of AAV recovered variants from in vivo screens. PacBio and SeqCenter for PacBio sequencing of iPSC recovered variants. UT Southwestern Medical Center’s Center for Alzheimers’ and Neurodegenerative Diseases, Dr. Marc Diamond and Dr. Clarissa Bhanvadia for iPSC cortical neuron differentiation and cultures. The Peter O’Donnell Jr. Brain Institute and Dr. Allan-Hermann Pool’s MERSCOPE Core, UTSW’s Cryo-EM core facilities, UTSW Whole brain microscopy core facilities. This project was supported by the following funding sources: NIH/NINDS awards P51OD011104 (Hamm), U42OD024282 (Russell-Lodrigue), U42OD010568 (Russell-Lodrigue), and R01NS095867 (to S.J.G), as well as support from the Haggerty Foundation (to S.J.G.), the UTSW Peter O’Donnell Jr. Brain Institute Neural Scientist Training Program Fellowship (to I.T.G), PacBio STAR Grant (to I.T.G), UTSW Peter O’Donnell Jr. Brain Institute Sprouts Grant (to I.T.G.), and UTSW Mechanisms of Disease and Translational Sciences NIH T32 Training Grant GM109776 (to I.T.G).

## Author contributions

I.T.G and S.J.G conceived and designed the study. E.A.L. designed the controlled mutagenesis AAV9 library, Y.H. processed tissues and tissue staining, I.T.G collected and analyzed histology images, I.T.G. performed and analyzed MERFISH experiments, V.R. prepared AAV libraries, recovered AAV genomes from tissues and processed samples for biodistribution, I.T.G. produced recombinant AAV for in vivo experiments, M.R. performed mouse injections, M.T. produced AAV9 controlled randomization library, recovered AAV genomes from tissues, and processed samples for SMRT sequencing, I.T.G. and H.H. performed manufacturability, infectivity assays, binding assays, produced recombinant AAV for in-vitro experiments, zeta potential, and performed viral stability assays, R.A. performed multiplex reporter QC experiments, I.T.G., P.K., and D.K. performed Cryo-EM collection and processing. carried out sample preparation, data analysis, manuscript writing, interpreted data and conceptualization of study. I.T.G. and L.A.J. analyzed and interpreted computational Rosetta predictions and Cryo-EM structure, I.T.G., M.R., S.J.G, and R.M.B. designed AAV circulation kinetics experiment, I.T.G., A.K. and C.X. carried out data processing of SMRT sequencing and data analysis, C.B.T. and S.E.B. carried out NHP injections and NHP sample collection, I.T.G. wrote the manuscript and prepared the figures. S.J.G. wrote and edited the manuscript. S.J.G. supervised all aspects of the work. All authors read and approved the final manuscript.

## REFERENCES

1 Ling, Q., Herstine, J. A., Bradbury, A. & Gray, S. J. AAV-based in vivo gene therapy for neurological disorders. Nature Reviews Drug Discovery 22, 789–806 (2023). 10.1038/s41573-023-00766-7

2 Dowling, J. J. et al. AAV gene therapy for hereditary spastic paraplegia type 50: a phase 1 trial in a single patient. Nat Med 30, 1882–1887 (2024). 10.1038/s41591-024-03078-4

3 Xie, J. et al. Effective and Accurate Gene Silencing by a Recombinant AAV-Compatible MicroRNA Scaffold. Mol Ther 28, 422–430 (2020). 10.1016/j.ymthe.2019.11.018

4 Harper, S. Q. et al. RNA interference improves motor and neuropathological abnormalities in a Huntington’s disease mouse model. Proc Natl Acad Sci U S A 102, 5820–5825 (2005). 10.1073/pnas.0501507102

5 DeVos, S. L. et al. Tau reduction prevents neuronal loss and reverses pathological tau deposition and seeding in mice with tauopathy. Sci Transl Med 9 (2017). 10.1126/scitranslmed.aag0481

6 Amado, D. A. et al. AAV-based delivery of RNAi targeting Ataxin-2 improves survival, strength, and pathology in mouse models of rapidly and slowly progressive sporadic ALS. bioRxiv (2024). 10.1101/2024.01.31.578314

7 An, M. et al. In vivo base editing extends lifespan of a humanized mouse model of prion disease. Nat Med 31, 1319–1328 (2025). 10.1038/s41591-024-03466-w

8 Muller, A. et al. High-efficiency base editing in the retina in primates and human tissues. Nat Med 31, 490–501 (2025). 10.1038/s41591-024-03422-8

9 Chai, A. C. et al. Base editing correction of hypertrophic cardiomyopathy in human cardiomyocytes and humanized mice. Nat Med 29, 401–411 (2023). 10.1038/s41591-022-02176-5

10 DeFrancesco, L. Precision gene editing medicine makes history, and it’s just getting started. Nat Biotechnol 43, 1019–1022 (2025). 10.1038/s41587-025-02741-6

11 Bharucha-Goebel, D. X. et al. Intrathecal Gene Therapy for Giant Axonal Neuropathy. N Engl J Med 390, 1092–1104 (2024). 10.1056/NEJMoa2307952

12 Pian, J. M. Y. et al. How to pay for individualized genetic medicines. Nat Med 30, 1816–1818 (2024). 10.1038/s41591-024-03071-x

13 Mackall, C. L. et al. Enhancing pediatric access to cell and gene therapies. Nat Med 30, 1836–1846 (2024). 10.1038/s41591-024-03035-1

14 Mendell, J. R. et al. AAV gene therapy for Duchenne muscular dystrophy: the EMBARK phase 3 randomized trial. Nat Med 31, 332–341 (2025). 10.1038/s41591-024-03304-z

15 Qi, J. et al. AAV gene therapy for autosomal recessive deafness 9: a single-arm trial. Nat Med (2025). 10.1038/s41591-025-03773-w

16 Spuler, S., Borasio, G. D. & Grittner, U. Lessons from a negative gene therapy trial for Duchenne muscular dystrophy. Nat Med 31, 37–38 (2025). 10.1038/s41591-024-03316-9

17 Duan, D. Lethal immunotoxicity in high-dose systemic AAV therapy. Mol Ther 31, 3123–3126 (2023). 10.1016/j.ymthe.2023.10.015

18 Lee, Y., Jeong, M., Park, J., Jung, H. & Lee, H. Immunogenicity of lipid nanoparticles and its impact on the efficacy of mRNA vaccines and therapeutics. Exp Mol Med 55, 2085–2096 (2023). 10.1038/s12276-023-01086-x

19 Lauffer, M. C., van Roon-Mom, W., Aartsma-Rus, A. & Collaborative, N. Possibilities and limitations of antisense oligonucleotide therapies for the treatment of monogenic disorders. Commun Med (Lond) 4, 6 (2024). 10.1038/s43856-023-00419-1

20 Riedmayr, L. M. et al. mRNA trans-splicing dual AAV vectors for (epi)genome editing and gene therapy. Nat Commun 14, 6578 (2023). 10.1038/s41467-023-42386-0

21 Koblan, L. W. et al. In vivo base editing rescues Hutchinson-Gilford progeria syndrome in mice. Nature 589, 608–614 (2021). 10.1038/s41586-020-03086-7

22 Zhang, Y., Bassel-Duby, R. & Olson, E. N. CRISPR-Cas9 Correction of Duchenne Muscular Dystrophy in Mice by a Self-Complementary AAV Delivery System. Methods Mol Biol 2587, 411–425 (2023). 10.1007/978-1-0716-2772-3_21

23 Davis, J. R. et al. Efficient prime editing in mouse brain, liver and heart with dual AAVs. Nat Biotechnol 42, 253–264 (2024). 10.1038/s41587-023-01758-z

24 Byrne, B. J. et al. Current clinical applications of AAV-mediated gene therapy. Mol Ther 33, 2479–2516 (2025). 10.1016/j.ymthe.2025.04.045

25 Pupo, A. et al. AAV vectors: The Rubik’s cube of human gene therapy. Mol Ther 30, 3515–3541 (2022). 10.1016/j.ymthe.2022.09.015

26 Bryant, D. H. et al. Deep diversification of an AAV capsid protein by machine learning. Nat Biotechnol 39, 691–696 (2021). 10.1038/s41587-020-00793-4

27 Huang, Q. et al. Targeting AAV vectors to the central nervous system by engineering capsid-receptor interactions that enable crossing of the blood-brain barrier. PLoS Biol 21, e3002112 (2023). 10.1371/journal.pbio.3002112

28 Huang, Q. et al. An AAV capsid reprogrammed to bind human transferrin receptor mediates brain-wide gene delivery. Science 384, 1220–1227 (2024). 10.1126/science.adm8386

29 Tabebordbar, M. et al. Directed evolution of a family of AAV capsid variants enabling potent muscle-directed gene delivery across species. Cell 184, 4919– 4938 e4922 (2021). 10.1016/j.cell.2021.08.028

30 Zinn, E. et al. Ancestral library identifies conserved reprogrammable liver motif on AAV capsid. Cell Rep Med 3, 100803 (2022). 10.1016/j.xcrm.2022.100803

31 Deverman, B. E. et al. Cre-dependent selection yields AAV variants for widespread gene transfer to the adult brain. Nat Biotechnol 34, 204–209 (2016). 10.1038/nbt.3440

32 Goertsen, D. et al. AAV capsid variants with brain-wide transgene expression and decreased liver targeting after intravenous delivery in mouse and marmoset. Nat Neurosci 25, 106–115 (2022). 10.1038/s41593-021-00969-4

33 Leib, D. E. et al. Optimized AAV capsids for basal ganglia diseases show robust potency and distribution. Nat Commun 16, 4653 (2025). 10.1038/s41467-025-60000-3

34 Marx, V. Method of the year: long-read sequencing. Nat Methods 20, 6–11 (2023). 10.1038/s41592-022-01730-w

35 Casy, W. et al. SMRT Sequencing Enables High-Throughput Identification of Novel AAVs from Capsid Shuffling and Directed Evolution. Genes (Basel) 14 (2023). 10.3390/genes14081660

36 Scott, S. et al. AAVolve: Concatenated long-read deep sequencing enables whole capsid tracking during shuffled AAV library selection. Mol Ther Methods Clin Dev 32, 101351 (2024). 10.1016/j.omtm.2024.101351

37 Sonntag, F., Schmidt, K. & Kleinschmidt, J. A. A viral assembly factor promotes AAV2 capsid formation in the nucleolus. Proc Natl Acad Sci U S A 107, 10220– 10225 (2010). 10.1073/pnas.1001673107

38 Elmore, Z. C. et al. The membrane associated accessory protein is an adeno-associated viral egress factor. Nat Commun 12, 6239 (2021). 10.1038/s41467-021-26485-4

39 Bennett, A. et al. Thermal Stability as a Determinant of AAV Serotype Identity. Mol Ther Methods Clin Dev 6, 171–182 (2017). 10.1016/j.omtm.2017.07.003

40 Hsu, H. L. et al. Structural characterization of a novel human adeno-associated virus capsid with neurotropic properties. Nat Commun 11, 3279 (2020). 10.1038/s41467-020-17047-1

41 Moyer, T. C. et al. Highly conserved brain vascular receptor ALPL mediates transport of engineered AAV vectors across the blood-brain barrier. Mol Ther 33, 3902–3916 (2025). 10.1016/j.ymthe.2025.04.046

42 Ravindra Kumar, S., et al. Multiplexed Cre-dependent selection yields systemic AAVs for targeting distinct brain cell types. Nat Methods 17, 541–550 (2020). 10.1038/s41592-020-0799-7

43 Weinmann, J. et al. Identification of a myotropic AAV by massively parallel in vivo evaluation of barcoded capsid variants. Nat Commun 11, 5432 (2020). 10.1038/s41467-020-19230-w

44 DiMattia, M. A. et al. Structural insight into the unique properties of adeno-associated virus serotype 9. J Virol 86, 6947–6958 (2012). 10.1128/JVI.07232-11

45 Sonntag, F. et al. The assembly-activating protein promotes capsid assembly of different adeno-associated virus serotypes. J Virol 85, 12686–12697 (2011). 10.1128/JVI.05359-11

46 Hordeaux, J. et al. The Neurotropic Properties of AAV-PHP.B Are Limited to C57BL/6J Mice. Molecular Therapy 26, 664–668 (2018). 10.1016/j.ymthe.2018.01.018

47 Gonzalez, T. J. et al. Cross-species evolution of a highly potent AAV variant for therapeutic gene transfer and genome editing. Nature Communications 13, 5947 (2022). 10.1038/s41467-022-33745-4

48 Penzes, J. J. et al. Adeno-associated Virus 9 Structural Rearrangements Induced by Endosomal Trafficking pH and Glycan Attachment. J Virol 95, e0084321 (2021). 10.1128/jvi.00843-21

49 Tan, Y. Z. et al. Sub-2 A Ewald curvature corrected structure of an AAV2 capsid variant. Nat Commun 9, 3628 (2018). 10.1038/s41467-018-06076-6

50 Drouin, L. M. et al. Cryo-electron Microscopy Reconstruction and Stability Studies of the Wild Type and the R432A Variant of Adeno-associated Virus Type 2 Reveal that Capsid Structural Stability Is a Major Factor in Genome Packaging. J Virol 90, 8542–8551 (2016). 10.1128/JVI.00575-16

51 Bennett, A. et al. Structural and Biophysical Analysis of Adeno-Associated Virus Serotype 2 Capsid Assembly Variants. J Virol 97, e0177222 (2023). 10.1128/jvi.01772-22

52 Chen, X. et al. AAV9/MFSD8 gene therapy is effective in preclinical models of neuronal ceroid lipofuscinosis type 7 disease. J Clin Invest 132 (2022). 10.1172/JCI146286

53 Bailey, R. M., Armao, D., Nagabhushan Kalburgi, S. & Gray, S. J. Development of Intrathecal AAV9 Gene Therapy for Giant Axonal Neuropathy. Mol Ther Methods Clin Dev 9, 160–171 (2018). 10.1016/j.omtm.2018.02.005

54 Gray, S. J. et al. Production of recombinant adeno-associated viral vectors and use in in vitro and in vivo administration. Curr Protoc Neurosci **Chapter** 4, Unit 4 17 (2011). 10.1002/0471142301.ns0417s57

55 He, S. & Scheres, S. H. W. Helical reconstruction in RELION. J Struct Biol 198, 163–176 (2017). 10.1016/j.jsb.2017.02.003

56 Rohou, A. & Grigorieff, N. CTFFIND4: Fast and accurate defocus estimation from electron micrographs. J Struct Biol 192, 216–221 (2015). 10.1016/j.jsb.2015.08.008

57 Bepler, T. et al. Positive-unlabeled convolutional neural networks for particle picking in cryo-electron micrographs. Nat Methods 16, 1153–1160 (2019). 10.1038/s41592-019-0575-8

58 Pettersen, E. F. et al. UCSF Chimera--a visualization system for exploratory research and analysis. J Comput Chem 25, 1605–1612 (2004). 10.1002/jcc.20084

59 Goddard, T. D. et al. UCSF ChimeraX: Meeting modern challenges in visualization and analysis. Protein Sci 27, 14–25 (2018). 10.1002/pro.3235

60 Pettersen, E. F. et al. UCSF ChimeraX: Structure visualization for researchers, educators, and developers. Protein Sci 30, 70–82 (2021). 10.1002/pro.3943

61 Adams, P. D. et al. PHENIX: a comprehensive Python-based system for macromolecular structure solution. Acta Crystallogr D Biol Crystallogr 66, 213– 221 (2010). 10.1107/S0907444909052925

62 Afonine, P. V. et al. Real-space refinement in PHENIX for cryo-EM and crystallography. Acta Crystallogr D Struct Biol 74, 531–544 (2018). 10.1107/S2059798318006551

63 Chen, V. B. et al. MolProbity: all-atom structure validation for macromolecular crystallography. Acta Crystallogr D Biol Crystallogr 66, 12–21 (2010). 10.1107/S0907444909042073

64 Pachitariu, M. & Stringer, C. Cellpose 2.0: how to train your own model. Nat Methods 19, 1634–1641 (2022). 10.1038/s41592-022-01663-4

65 Palla, G. et al. Squidpy: a scalable framework for spatial omics analysis. Nat Methods 19, 171–178 (2022). 10.1038/s41592-021-01358-2

66 Wolf, F. A., Angerer, P. & Theis, F. J. SCANPY: large-scale single-cell gene expression data analysis. Genome Biol 19, 15 (2018). 10.1186/s13059-017-1382-0

