## Supplemental 1 for "A minimally modified AAV9 variant (AAV9fyhr) for enhanced CNS gene delivery by intra-CSF administration"

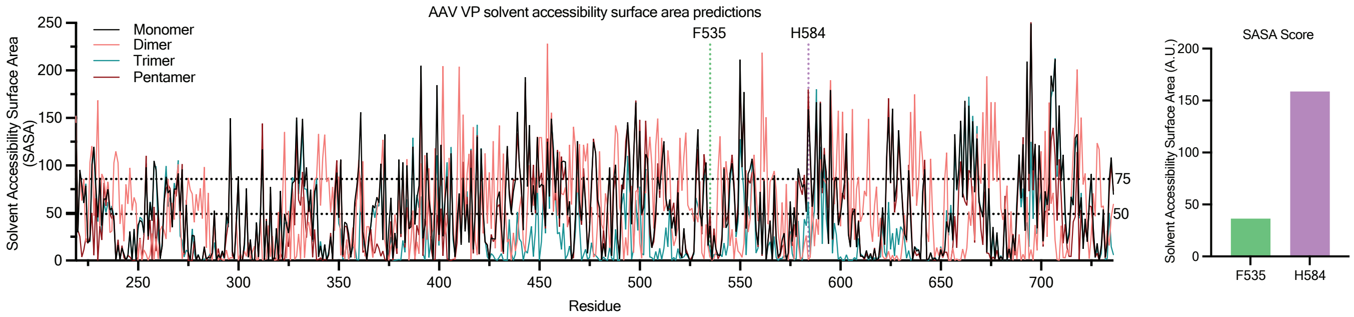


**Supplemental Fig. 1: in silico analysis of AAV9 viral protein solvent accessibility.** The viral protein of AAV9 (PDB: 7MT0) was analyzed using solvent accessibility surface area (SASA) per residue for the monomer (black), dimer (pink - two-fold), trimer (green - three-fold), and pentamer (purple - five-fold) states of capsid assembly. Position 535 (dashed green vertical line) and 584 (dashed purple vertical line). Histoplot displays the sasa score of 535 and 584 of the monomer state.
