## Supplemental 2 for "A minimally modified AAV9 variant (AAV9fyhr) for enhanced CNS gene delivery by intra-CSF administration"

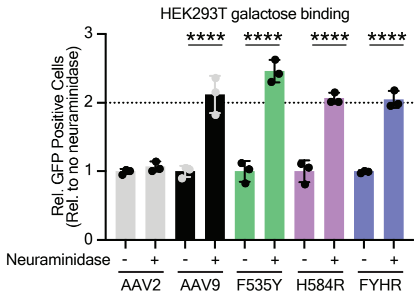


**Supplemental Fig. 2: AAV transduced neuraminidase treated HEK293T cells.** AAV capsid variants were used to transduce HEK293T cells with or without neuraminidase treatment. Cells were flow sorted for GFP positivity and compared relative to the HEK293T condition without neuraminidase treatment. One-way ANOVA was used with Sidak multiple comparison correction for transduction comparisons. Error bars indicate ± SD. ****p<0.0001
