## Supplemental 3 for "A minimally modified AAV9 variant (AAV9fyhr) for enhanced CNS gene delivery by intra-CSF administration"

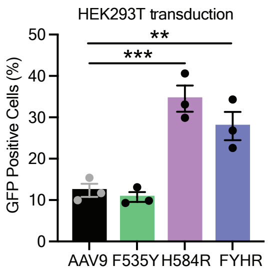


**Supplemental Fig. 3: HEK293T Transduction data.** Histoplots display flow cytometry quantification of GFP positive cells transduced with AAV capsid variants at 1 x 104 vg/cell after 48 hours post treatment. One-way ANOVA was used with Sidak multiple comparison correction for transduction comparisons. Error bars indicate ± SD. **$p<0.01$, ***$p<0.001$
