## Supplemental 4 for "A minimally modified AAV9 variant (AAV9fyhr) for enhanced CNS gene delivery by intra-CSF administration"

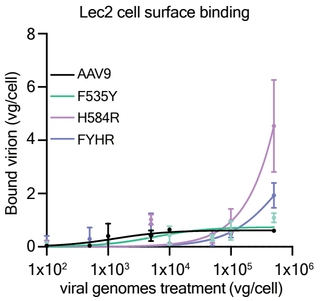


**Supplemental Fig. 4: Cell surface binding.** CHO lec2 cells were treated with increasing amounts of viral genomes per cell for each AAV variant. Binding of capsid particles were subsequently measured using quantitative PCR for Dnase resistant encapsidated genomes for viral particles bound to the cell surface.
