## Supplemental 5 for "A minimally modified AAV9 variant (AAV9fyhr) for enhanced CNS gene delivery by intra-CSF administration"

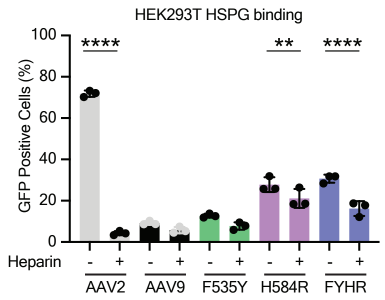


**Supplemental Fig. 5: Heparin competition assay in HEK293T cells.** Histoplots display GFP positive cell quantification of flow sorted HEK293T cells with and without heparin added to the AAV treatment. One-way ANOVA was used with Sidak multiple comparison correction for transduction comparisons. Error bars indicate ± SD. $^{**}p<0.01$, $^{****}p<0.0001$
