## Supplemental 6 for "A minimally modified AAV9 variant (AAV9fyhr) for enhanced CNS gene delivery by intra-CSF administration"

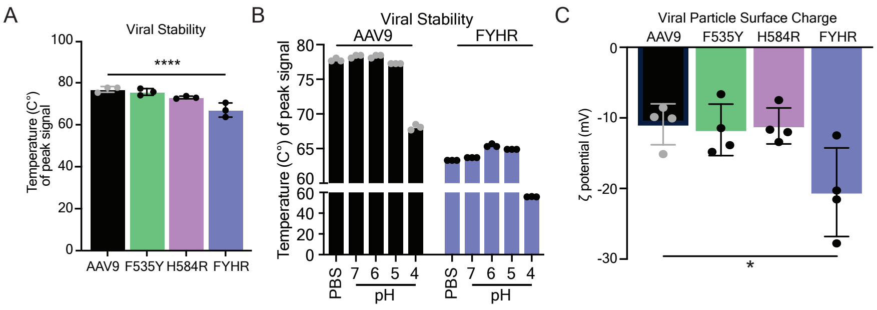


**Supplemental Fig. 6: Biophysical properties of AAV9 variants measuring viral stability and particle surface charge.** (A) Histoplot displays thermal stability of viral capsids at physiological pH. n=3 per condition (B) Demonstrates the viral stability at pH 7 (PBS and sodium bicarbonate buffer), 6, 5, and 4 (sodium bicarbonate buffer). n=3 per condition (C) Zeta potential measurements of particle charge are displayed in histoplots. n=4 per variant. One-way ANOVA was used with Sidak multiple comparison correction for transduction comparisons. Error bars indicate ± SD. $*p<0.05$, $****p<0.0001$
