## Supplemental 7 for "A minimally modified AAV9 variant (AAV9fyhr) for enhanced CNS gene delivery by intra-CSF administration"

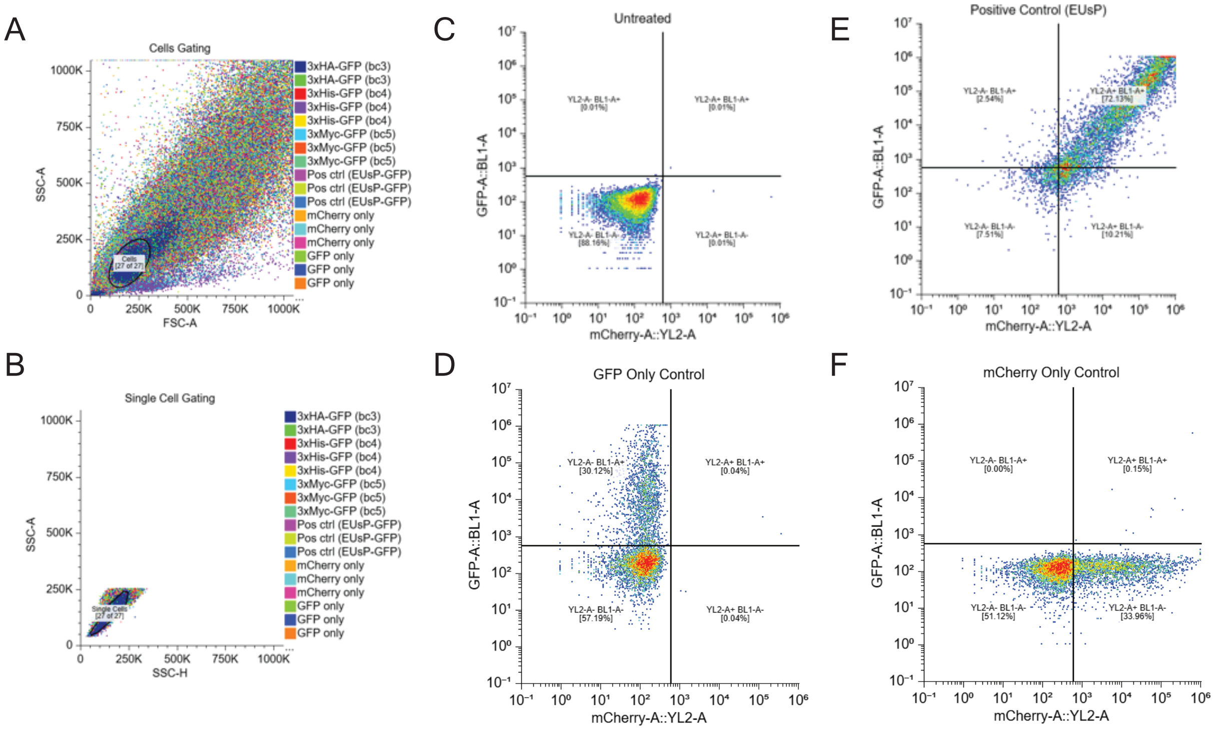


**Supplemental Fig. 7: Reporter vector validation.** (A-F) Gating of flow cytometry quantification of multiplexed reporter vectors displaying cells (A), single cells (B), non-transfected cell sample (C), GFP single transfection control (D), positive control sample (E), and mCherry control single transfection (F).
