## Supplemental 8 for "A minimally modified AAV9 variant (AAV9fyhr) for enhanced CNS gene delivery by intra-CSF administration"

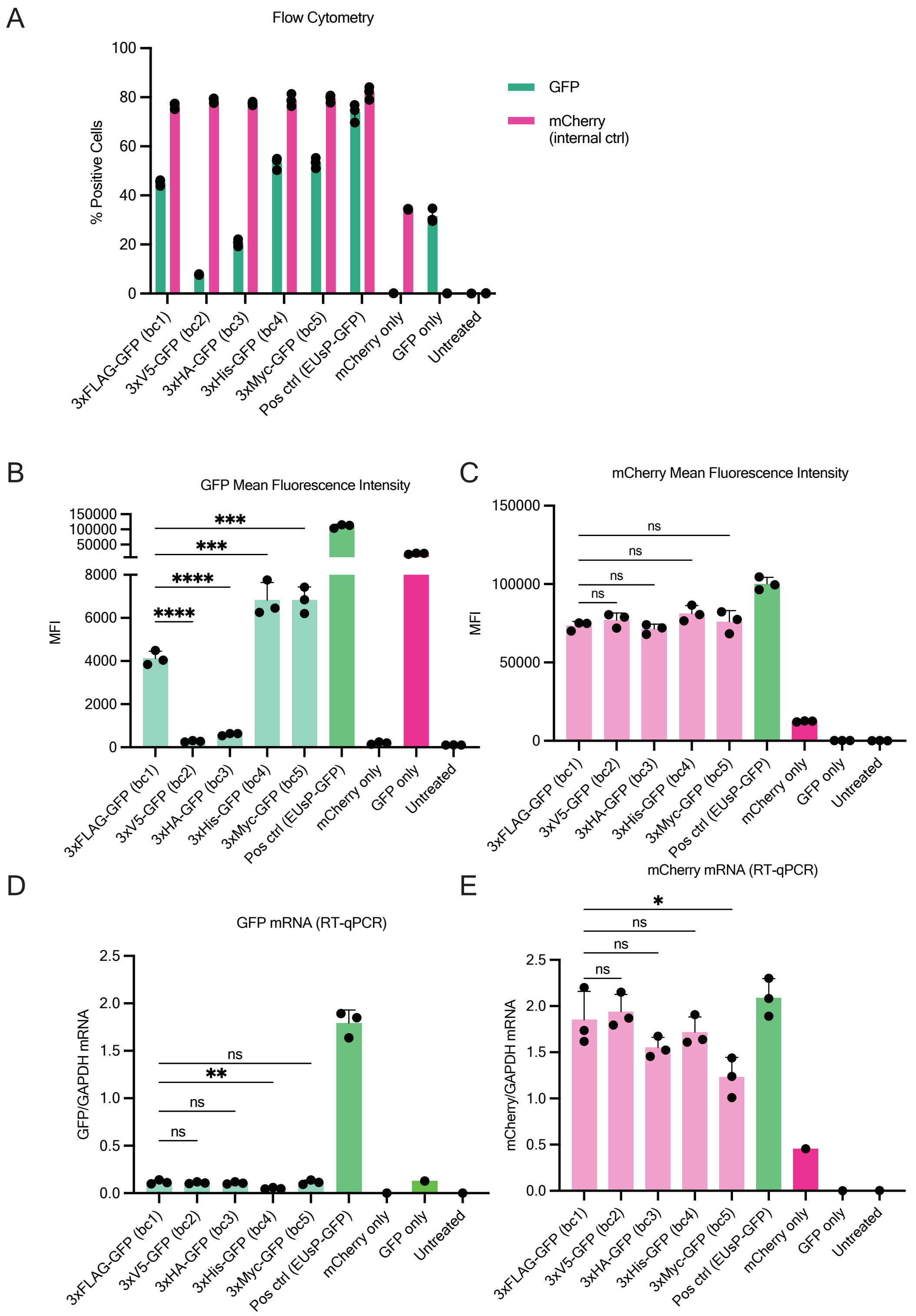


**Supplemental Fig. 8: Reporter vector validation quantification.** (A-E) Quantification of protein and RNA expression of multiplex reporter vectors. (A) Histoplots display flow cytometry quantification of GFP and mCherry positive cells transfected with reporter vectors or control expression plasmids. (B & C) Bar plots display the mean fluoresence intensity of GFP (B) and mCherry (C) positive cells. (D & E) Histoplots display relative mRNA expression of GFP (D) and mCherry (E). One-way ANOVA was used with Dunnett multiple comparison correction for transduction comparisons. Error bars indicate ± SD. *p<0.05, **p<0.01, ***p<0.001, ****p<0.0001
