## Supplementary figures and images for "A minimally modified AAV9 variant (AAV9fyhr) for enhanced CNS gene delivery by intra-CSF administration"

### Supplemental 9

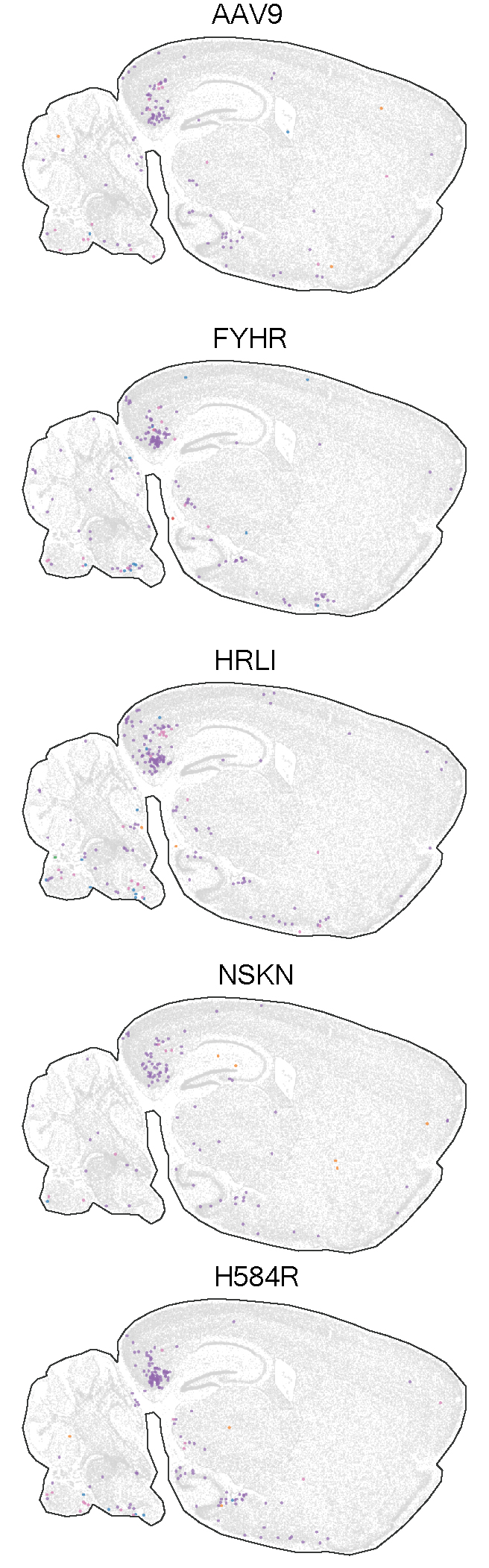
