## Supplemental 10 for "A minimally modified AAV9 variant (AAV9fyhr) for enhanced CNS gene delivery by intra-CSF administration"

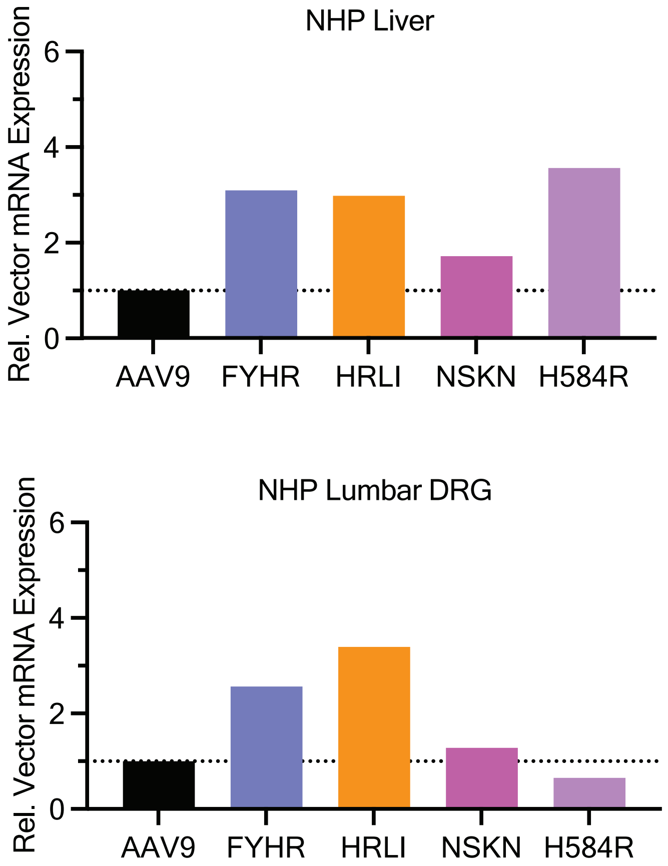


**Supplemental Fig. 10: NHP Liver and lumbar dorsal root ganglion (DRG) RNAScope.** AAV vector expression in nonhuman primate liver. Histoplot displays relative mRNA expression (rel. to AAV9) of each capsid variant.
