## Supplemental 11 for "A minimally modified AAV9 variant (AAV9fyhr) for enhanced CNS gene delivery by intra-CSF administration"

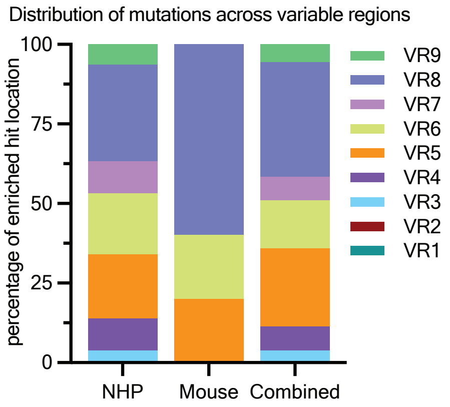


**Supplemental Fig. 11: AAV9 variant hit hotspots.** Variants recovered from screens in mice and nonhuman primates. stacked histoplots displays the proportion of hits recovered from mice, NHPs, and combined across the various variable regions (VRs).
