## Supplemental 12 for "A minimally modified AAV9 variant (AAV9fyhr) for enhanced CNS gene delivery by intra-CSF administration"

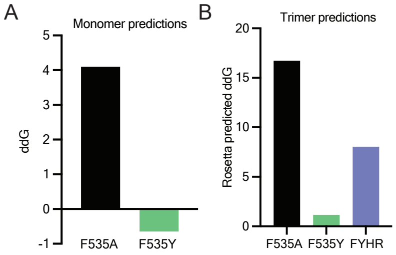


**Supplemental Fig. 12: Rosetta in silico ddG predictions.** (A) Prediction of position 535 to an alanine or tyrosine in the context of the VP3 monomer. (B) ddG predictions of position 535 to alanine, tyrosine, or double mutant FYHR in the context of the trimer (three-fold axis of symmetry).
