## Supplemental 13 for "A minimally modified AAV9 variant (AAV9fyhr) for enhanced CNS gene delivery by intra-CSF administration"

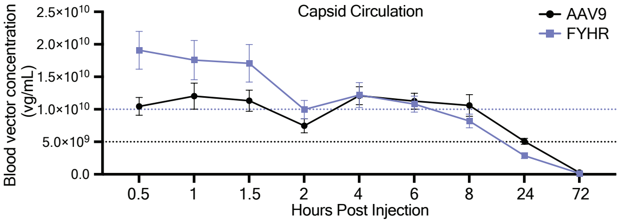


**Supplemental Fig. 13:. Capsid circulation kinetics in mice.** Blood collected from mice at various timepoints after CSF intrathecal lumbar cistern administration of AAV9 and FYHR was probed for Dnase resistant encapsidated viral genomes using quantitative PCR. Dotted lines represent EC50 of each capsid variant. (n=12; 6M;6F)
