## Supplemental Methods and Supplemental References for "A minimally modified AAV9 variant (AAV9fyhr) for enhanced CNS gene delivery by intra-CSF administration"

Multiplex reporter validation

HEK293T cells were seeded onto 48-well plates. 24 hours post seeding, cells were transfected with expression plasmids at equimolar concentrations using lipofectamine 3000 (Thermo) per manufacturer's specifications. 48 hours post transfection, cells were collected. For flow cytometry quantification, cells were sorted for GFP and mCherry. Adjustments for fluorophore positive signal was adjusted to single plasmid transfection controls and untreated to minimize any background signal. For mRNA quantification, RNA was extracted using RNeasy Plus kit (Qiagen) per manufacturer's specification. cDNA was synthesized from RNA using Superscript IV VILO plus ezDNase kit (Invitrogen). qPCR was amplified using primers targeting GFP (forward: 5'-AGTCCGCCCTGAGCAAAGA-3', reverse: 5'-TCCAGCAGGACCATGTGATC-3'), mCherry (forward: 5'-agggcgaggaggataacatg-3', reverse: 5'-cttggtcaccttcagcttgg-3'), and GAPDH (forward: 5'-TTGGCTACAGCAACAGGGTG-3', reverse: 5'-GGGGAGATTCAGTGTGGTGG-3') with PowerUp SYBR Green Master Mix (Applied Biosciences). Relative quantification was performed using the $2^{-dCt}$ method of fluorophore expression relative to GAPDH. Data was analyzed using one-way ANOVA with comparison to bc1 using Dunnett's multiple comparisons test.

Viral Transduction Assays

Cells (HEK293T, Lec2, SH-SY5Y, Neuro-N2A, Cos7, and iPSC-derived cortical neurons) were seeded into 96-well flat bottom plates with 10,000 cells per well. 24 hours after seeding, cells were treated with $1 \times 10^4$, $1 \times 10^5$, or $1 \times 10^6$ vg per cell. Cells were collected 48 hours after virus treatment for immortalized cells, or seven days post transduction for iPSC derived cortical neurons. In brief, growth medium was removed, cells were gently washed with 1X phosphate buffered saline, dissociated with 0.25% trypsin, neutralized with outgrowth medium, and then transferred to a round bottom 96-well plate before pelleting by centrifugation. Medium was removed and cells were fixed with 4% PFA at room temperature for 15 minutes. PFA was removed and cells were resuspended using 1X PBS. Cells were quantified for GFP+ cells using an Attune Cytpix (Thermo) flow cytometer.

Specifically for enzymatic desialylation experiments, the methods were adapted and modified as previously described[1]. Briefly, HEK293T cells were seeded into 96-well flat bottom plates with 10,000 cells per well. 24 hours after seeding, cells were treated with Neuraminidase [50 milliunits/mL] (Vibrio cholerae; Sigma N7885) or Heparinase I [3 units/mL] & III [1.5 units/mL] (Flavobacterium heparinum; Sigma H2519 and H8891) for 4 hours at 37 °C followed by $1 \times 10^5$ vg per cell. Cells were collected 24 hours after virus treatment.

Specifically for the heparin competition assay, HEK293T cells were seeded into 96-well flat bottom plates with 10,000 cells per well. 24 hours after seeding, cells were treated with $1 \times 10^5$ vg per cell supplemented with heparin [1,000ug/mL] (Sigma), adapted and modified as previously described[2]. Cells were collected 24 hours after virus treatment.

Zeta potential

AAV was diluted to $1.0 \times 10^9$ vg/mL and zeta potential was measured using a NanoBrook system (Brookhaven Instruments). 500 µL of sample was added into the sample cuvette and PALS was used to determine zeta potential. Three measurements were recorded for each sample.

Differential scanning fluorimetry

Viral stability of each capsid variant was characterized and adapted as previously described[3]. In brief, virus samples were incubated in PBS or Sodium Acetate Buffer [pH 7, 6, 5, or 4] with 1% SYPRO orange dye total of 50 uL reaction volumes. Measurements were collected on a QuantStudio 6 Pro with temperature ranging from 30 °C to 99 °C and ramping at 0.5 °C per step.

In silico rosetta ddG predictions

The AAV9 VP structure (PDB: 3UX1) was prepared for in silico estimation of assembly energies. PyMol was used to prepare the monomeric, dimeric, trimeric and pentameric assemblies of AAV9 and used as input for subsequent mutagenesis and minimization using RosettaScripts interface to Rosetta v3.12[4,5]. Changes in assembly energy were calculated as previously described using an adaptation of the Flex ddG protocol[6,7]. Solvent accessibility surface area, SASA, was performed by utilizing the prior assemblies. SASA was calculated in Rosetta using 'TotalSasa' mover with the 'report_per_residue_sasa' flag set to true[4]. To determine the interface of multimeric assemblies in the dimer, trimer, and pentamer, the difference of each assembly's SASA values were determined by subtracting the monomer SASA values.

Capsid circulation kinetics

Adult mice (n=12; 6M;6F) were injected via a lumbar intrathecal route with pooled AAV9 and FYHR at $2 \times 10^{11}$ vg total ($1 \times 10^{11}$ vg per capsid). Blood samples were collected by tail nick at 0.5, 1, 1.5, 2, 4, 6, 8, 24, and 72 hours post injection. Viral genomes were recovered from blood using DNeasy blood and tissue kit (Qiagen) and vector genomes were determined by qPCR targeting the ITGBC1 and ITGBC2 DNA barcodes and absolute quantification was determined with a linearized plasmid DNA standard curve.

Supplemental References


1 Shen, S., Bryant, K. D., Brown, S. M., Randell, S. H. & Asokan, A. Terminal N-linked galactose is the primary receptor for adeno-associated virus 9. *J Biol Chem* **286**, 13532–13540 (2011). https://doi.org/10.1074/jbc.M110.210922

2 Schmidt, M. *et al.* Adeno-associated virus type 12 (AAV12): a novel AAV serotype with sialic acid- and heparan sulfate proteoglycan-independent transduction activity. *J Virol* **82**, 1399–1406 (2008). https://doi.org/10.1128/JVI.02012-07

3 Bennett, A. *et al.* Thermal Stability as a Determinant of AAV Serotype Identity. *Mol Ther Methods Clin Dev* **6**, 171–182 (2017). https://doi.org/10.1016/j.omtm.2017.07.003

4 Fleishman, S. J. *et al.* RosettaScripts: a scripting language interface to the Rosetta macromolecular modeling suite. *PLoS One* **6**, e20161 (2011). https://doi.org/10.1371/journal.pone.0020161

5 Leman, J. K. *et al.* Macromolecular modeling and design in Rosetta: recent methods and frameworks. *Nat Methods* **17**, 665–680 (2020). https://doi.org/10.1038/s41592-020-0848-2

6 Barlow, K. A. *et al.* Flex ddG: Rosetta Ensemble-Based Estimation of Changes in Protein-Protein Binding Affinity upon Mutation. *J Phys Chem B* **122**, 5389–5399 (2018). https://doi.org/10.1021/acs.jpcb.7b11367

7 Mullapudi, V. *et al.* Network of hotspot interactions cluster tau amyloid folds. *Nat Commun* **14**, 895 (2023). https://doi.org/10.1038/s41467-023-36572-3
